# Direct β1/ β2 AMPK activation reduces liver steatosis but not fibrosis in a mouse model of non-alcoholic steatohepatitis

**DOI:** 10.1101/2024.05.30.596624

**Authors:** Karly M. Mather, Michelle L. Boland, Emma L. Rivers, Abhishek Srivastava, Marianne Schimpl, Paul Hemsley, James Robinson, Paul T. Wan, Josefine Hansen, Jon A. Read, James L. Trevaskis, David M. Smith

**Affiliations:** Bioscience, Research and Early Development, Cardiovascular, Renal and Metabolism (CVRM), BioPharmaceuticals R&D, AstraZeneca, Gaithersburg, MD USA; Hit Discovery, Discovery Sciences, R&D, AstraZeneca, Cambridge, UK; Drug Metabolism and Pharmacokinetics, Clinical Pharmacology and Safety Sciences, R&D, AstraZeneca, Cambridge, UK; Structure and Biophysics, Discovery Sciences, R&D, AstraZeneca, Cambridge, UK; Discovery Biology, Discovery Sciences, R&D, AstraZeneca, Cambridge, UK; Emerging Innovations Unit, Discovery Sciences, R&D, AstraZeneca, Cambridge, UK

**Keywords:** NASH, AMPK, NAFLD, Fibrosis, Steatosis

## Abstract

5’AMP-activated protein kinase (AMPK) activators show potential for treating Non-Alcoholic Fatty Liver Disease (NAFLD) and Non-Alcoholic Steatohepatitis (NASH) due to their inhibiting effects on fatty acid and cholesterol synthesis. The absence of treatments for NASH, and its propensity for progression to severe disease, lead us to identify and characterize BI9774, a small molecule AMPK activator, which we used to evaluate this potential, including its ability to reduce the NASH specific qualities of fibrosis and inflammation in a preclinical study.

Male *Lep^ob^/Lep^ob^* mice on a control or NASH inducing (AMLN) diet, with liver fibrosis were given BI9774 or vehicle for 6 weeks while metabolic and NASH endpoints were evaluated.

BI9774 treatment decreased plasma ALT, terminal liver weight, and liver lipids. RNA expression of collagen-related genes decreased, although collagen protein and inflammation remained unaltered. We also observed increased heart weight and glycogen levels, and increased expression of genes associated with cardiac hypertrophy.

AMPK activation improved many metabolic endpoints, but lack of significant improvement in liver fibrosis and negative cardiac effects suggest systemic AMPK activation is not an ideal NASH therapy. Reductions in steatosis and fibrosis-related genes indicate that, with extended treatment, a liver specific AMPK activator has potential to resolve hepatic fibrosis.

**Summary Statement:** Fatty liver disease affects up to 30 percent of adults worldwide with 30% of patients progressing to more sever liver disease. AMPK activation can help reduce liver fat.

## Introduction

Non-alcoholic fatty liver disease (NAFLD), typically defined by >5% hepatic steatosis, is the leading chronic liver disease worldwide, potentially affecting up to 25% of the adult population (Younossi et al., 2016). NAFLD is highly associated with metabolic diseases such as obesity, cardiovascular issues, and diabetes. In a recent study, two thirds of type 2 diabetic patients were shown to have evidence of NAFLD (Konerman et al., 2018). These patients can be asymptomatic and are often self-resolving, however, around 30% progress to develop non-alcoholic steatohepatitis (NASH) (Williams et al., 2011), which is characterized by inflammation. This inflammation often leads to fibrosis, cirrhosis, and hepatocellular carcinoma, which currently have no FDA approved therapies available.

5’AMP-activated protein kinase (AMPK) is a key regulator of energy balance, and AMPK activation has long been touted as a possible therapeutic strategy for the treatment of metabolic disorders (Grahame Hardie, 2014, Carling, 2017, Cool et al., 2006). Clinical development of AMPK activators has been hindered by safety challenges associated with direct kinase activation and other pharmacokinetic issues (Olivier et al., 2018). Nevertheless, recent pharmacological and genetic advances support a role for AMPK activators as a potential treatment for NAFLD/NASH due to its ability to directly inhibit acetyl CoA carboxylase (ACC) and stimulate malonyl CoA decarboxylase, thereby increasing fatty acid oxidation (Steinberg et al., 2006). AMPK activation also decreases cholesterol synthesis through phosphorylating and thereby inactivating the rate limiting enzyme HMG-CoA reductase (DeBose-Boyd, 2008, Soto-Acosta et al., 2017). Esquejo and colleagues showed the β1 AMPK activator PF-06409577 lowered liver lipid, as well as plasma triglycerides and cholesterol in mice, rats, and monkeys (Esquejo et al., 2018). Woods et al induced a liver-specific mutation in mice, which led to increased AMPK activation. These mice exhibited protection from high-fructose diet induced liver steatosis through inhibition of *de novo* lipogenesis but showed no change in fatty acid oxidation (Woods et al., 2017). Garcia et al created a mouse with an inducible activated AMPK alpha1 subunit directed to the liver. These mice were also protected from high fat diet-induced hepatic steatosis and showed decreased inflammation and fibrosis related gene expression (Garcia et al., 2019). Hu et al showed liver protection with activation of AMPK using 5-aminoimidazole-4-carboxamide-1-4-ribofuranoside (AICAR), which ameliorated portal hypertension and liver cirrhosis in rats (Hu et al., 2019).

Due to these promising effects shown by AMPK activation on metabolic disease, we desired to determine whether a systemic small molecule could induce these effects to the extent of mitigating the attributes of NAFLD and NASH. To do this, we characterized several publicly known AMPK activating compounds including the Boeringer Ingelheim compound BI9774 (example 2 in patent WO2015007669) (Himmelsbach F, 2015), (Boehringer-Ingelheim, 2021). Analysis of this compound in enzyme and cell-based systems showed the compound had appropriate drug metabolic and pharmacokinetic properties for a murine in-vivo study. We then tested the effect of BI9774 administration on liver-related endpoints in the high *trans*-fat/high fructose/high cholesterol fed leptin-deficient *Lep^ob^/Lep^ob^*mouse NASH model. This model exhibits several hallmark features of NASH (steatosis, inflammation, and fibrosis) with obesity and perturbed glycemic control (Trevaskis et al., 2012), (Hansen et al., 2017). Here we show that small molecule AMPK activation was effective at reducing liver weight, liver lipid and collagen gene expression but had minimal changes on fibrosis.

## Results

### BI9774 is a specific and selective AMPK activator with optimal properties for in-vivo studies

BI9774 showed equal potency agonist against both β1 and β2 beta-isoforms with EC50s of 20.0 nM and 63.1 nM respectively in isolated human enzyme assays. This is optimal because the target tissue for this study was mouse liver, which predominantly expresses β1 AMPK, while human liver is primarily β2. Phosphorylation of ACC by BI9774 in the human HepG2 hepatoma cell line showed very potent activation with an EC50 of 83.2 nM. The physical properties and *in vitro* metabolism of the compound are outlined in Table 1. Overall BI9774 showed good permeability, low protein binding, low hepatic metabolism, and moderate solubility, making it a good in-vivo candidate. After being tested against a panel of 408 protein kinases (S1 table), and in dose response against a panel of 70 other diverse targets (S2 Table), all targets showed at least 100-fold lower potency than the AMPK EC_50_ with no significant interactions. This allows us to reasonably conclude that the activity seen with BI9774 treatment is due to AMPK activation.

**Table 1.** Properties of BI9774. *^a^*EC50 values reported as the mean of 10 experiments for β1 and 5 experiments for β2 ± standard error of the mean (SEM). *^b^*EC50 value reported as the mean of 3 separate experiments ± SEM. *^c^*Distribution coefficient between 1-octanol and aqueous phosphate buffer at pH 7.4. *^d^*Solubility in aqueous phosphate buffer at pH 7.4 after 27 h at 25 °C. *^e^*Intrinsic permeability Papp using Caco2 cells. *^f^*Binding to human (h), rat (r) and mouse (m) plasma protein determined by equilibrium dialysis. *^g^*Total intrinsic clearance (CL_int_) in human microsomes. *^h^*Total intrinsic clearance (CLint) in human (h), rat (r) and mouse (m) hepatocytes.

| Property | Value |
| --- | --- |
| AMPK $\alpha 1\beta 1\gamma 1$ $EC_{50}$<sup>a</sup> | 20.0 nM ± 9.5 |
| AMPK $\alpha 1\beta 2\gamma 1$ $EC_{50}$<sup>a</sup> | 63.1 nM ± 14.7 |
| Hu HepG2 $EC_{50}$<sup>b</sup> | 83.2 nM ± 13.4 |
| MW / cLogP / tPSA | 541 / 3.8 / 122 |
| Log $D_{7.4}$<sup>c</sup> | 2.8 |
| Aq. Solubility (μM)<sup>d</sup> | 8 |
| Permeability ($1 \times 10^{-6}$ cm/s)<sup>e</sup> | 26 |
| Plasma Protein Binding: h / r / m (% free)<sup>f</sup> | 3.7 / 2.6 / 2.6 |
| Human Microsomes Clearance (μL/min/mg)<sup>g</sup> | <3 |
| Hepatocytes Clearance: h / r / m (μL/min/$10^6$ cells)<sup>h</sup> | <1 / <1.4 / 8.6 |

### BI9774 binds at the ADaM site

To confirm mechanism, a crystal structure of BI9774 bound to human AMPK (α2β1γ1) at 2.5 Å resolution was generated (Figure 1). This shows the compound binds at the ADaM site, similar to the other direct AMPK activators: 991 (Xiao et al., 2013), A-769662 (Calabrese et al., 2014), PF-739 (Myers et al., 2017), and MK-8722 (Cokorinos et al., 2017). The molecular interactions observed between BI9774 and AMPK offer a structural rationale for the ability of BI9774 to activate AMPK complexes containing either the β1 or β2 subunit and was used as validation of target engagement, PDB deposition ID 8BIK (Supplementary Figure 1).

**Fig 1.**
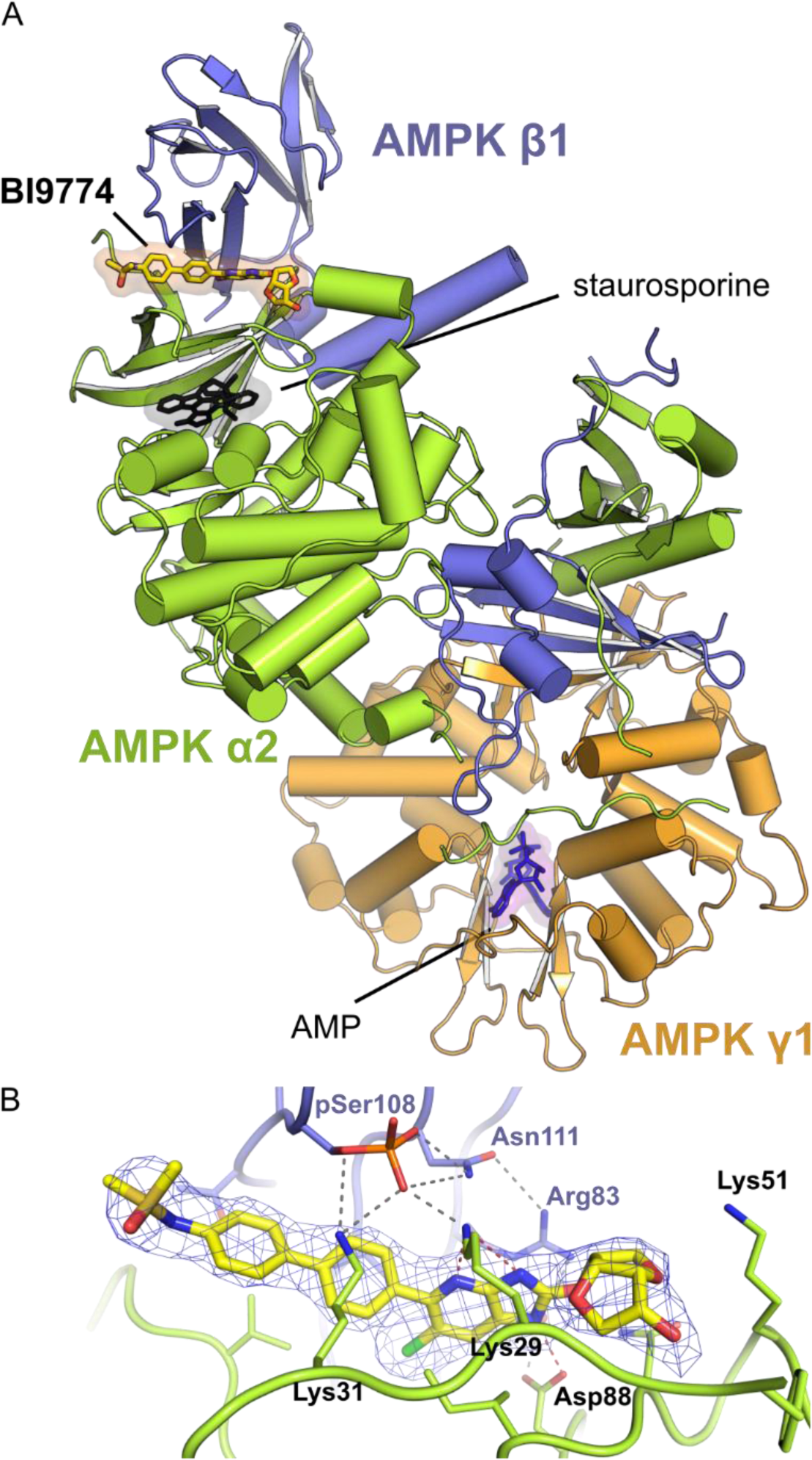
Crystal structure of AMPK in complex with allosteric activator BI9774. Activator BI9774 (yellow sticks) binds at the interface between the catalytic alpha-subunit (green) and the carbohydrate-binding module of the regulatory beta-subunit (blue) otherwise known as the ADaM site. The kinase active site is occupied by staurosporine (black sticks); and in the gamma-subunit (orange) 2 of the 3 AMP binding sites are occupied (purple sticks). **B**. Close-up view of the molecular interactions in the ADaM site. Multiple hydrogen bonds (dashed lines) between the alpha and beta subunit are dependent on phospho-Ser108 of the beta-subunit. Electron density (2F_o_-F_c_) for the ligand is shown as blue mesh, contoured at 1 σ.

### Pharmacokinetic data shows ideal bioavailability in Lep^ob^/Lep^ob^ mice

Plasma pharmacokinetics in C57BL6J mice demonstrated good oral bioavailability and half-life, while intravenous administration showed low clearance (Table 2). Single oral doses of 30 mg/kg and 3 mg/kg were tested for *in vivo* bioavailability in *Lep^ob^/Lep^ob^* mice, (Figure 2). The 30 mg/kg dose showed free drug concentrations above the isolated enzyme EC_50_ for over 18 hours whereas the 3 mg/kg dose showed exposure above the EC_50_ for just under 5 hours. This correlates with Boehringer Ingelheim’s own data showing liver target engagement at 3 mg/kg and maximal activity at 30 mg/kg (Boehringer-Ingelheim, 2021), confirming these are appropriate doses for our NASH *Lep^ob^/Lep^ob^* mouse study.

**Fig 2:**
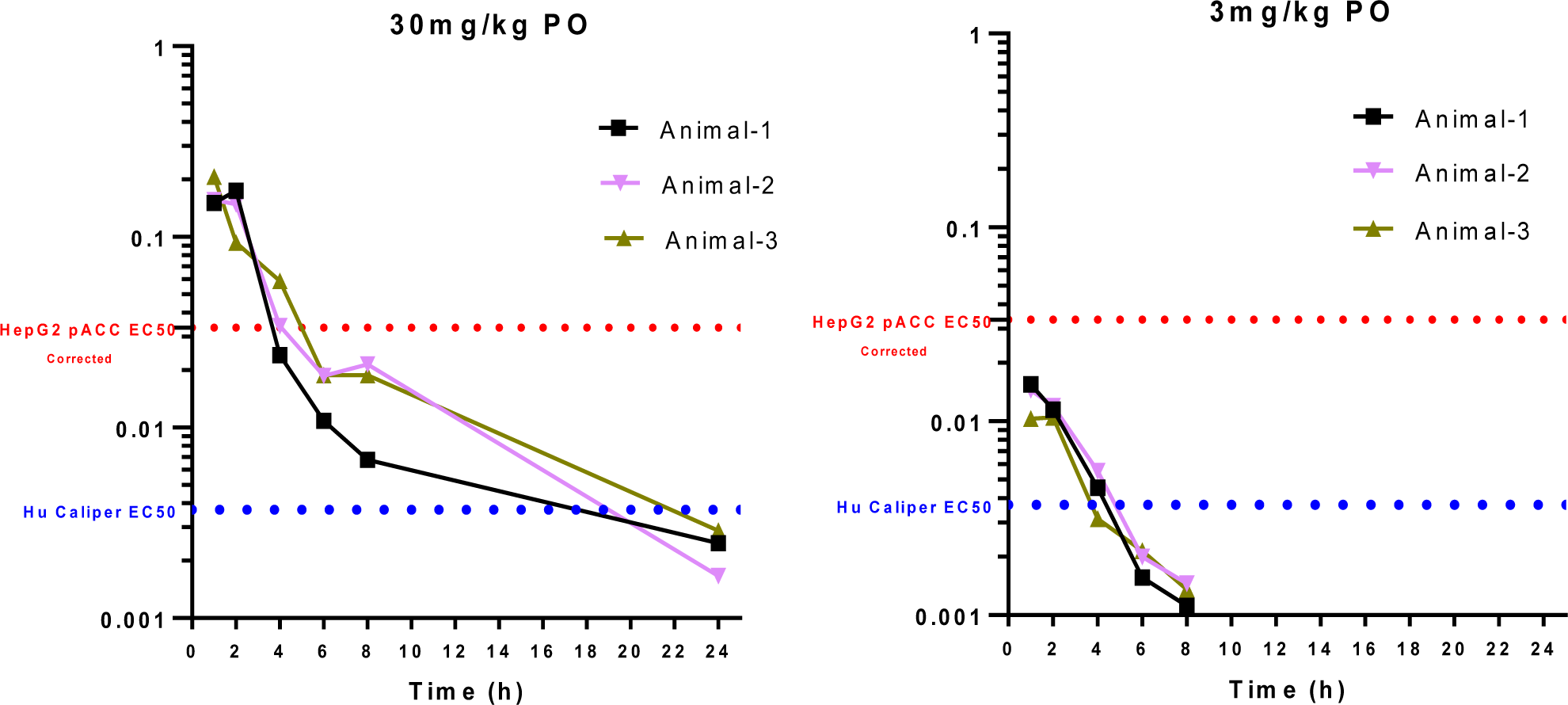
Pharmacokinetics of BI9774 in *Lep^ob^/Lep^ob^* mice (30, 3 mg/kg – free exposure) *Lep^ob^/Lep^ob^* mice (n=3 per group) were given a single dose of BI9774 by oral gavage of either 3 or 30 mg/kg body weight. The vertical axis shows the free concentration (μM) taking into account mouse plasma protein binding (2.6% free). The blue dotted line shows the EC_50_ for BI9774 in the recombinant enzyme assay. The red dotted line shows the EC_50_ for BI9774 in the HepG2 ACC phosphorylation assay corrected for free compound level. A single dose of 30 mg/kg resulted in free drug concentrations above the isolated enzyme EC_50_ for over 18 h whereas the lower dose of 3 mg/kg only showed exposure above the EC_50_ for less than 5h.

**Table 2:** Mouse in Vivo Pharmacokinetic Profile for BI9774. Male C57BL/6 mice were dosed with BI9774 either administered orally (5 mg/kg body weight) or intravenously (1 mg/kg body weight) as a solution containing 5% DMSO, 95% SBE-ß-CD (30% w/v) in water. Blood samples obtained postdosing were collected in tubes containing EDTA-K2 and centrifuged at 4000 × g for 5 min to obtain plasma. The concentration of BI9774 in plasma samples was determined using a liquid chromatography-tandem mass spectrometry (LC-MS/MS) method and pharmacokinetic parameters derived as in the methods. **^a^**Cmax is the maximum concentration (µM) obtained from a single oral dose of 5 mg/kg body weight. **^b^**AUC (μM.h) is the area under the concentration curve with time following oral and intravenous dosing. **^c^**T½(h) is the compound half-life. **^d^**Plasma clearance rate expressed in mL/min/kg. **^e^**Volume of distribution at steady state in L/kg. **^f^**Oral bioavailability (%).

| Plasma pharmacokinetics in C57BL6 mouse | Intravenous (1 mg/kg) | Oral (5 mg/kg) |
| --- | --- | --- |
| C<sub>max</sub> (μM)<sup>a</sup> | | 1.54 |
| AUC (μM*h)<sup>b</sup> | 2.3 | 5.12 |
| T<sub>1/2</sub> (h)<sup>c</sup> | 2.8 | 3 |
| Clearance (ml/min/kg)<sup>d</sup> | 13.5 | - |
| V<sub>ss</sub> (L/kg)<sup>e</sup> | 0.78 | - |
| Bioavailability (%)<sup>f</sup> | - | 45 |

### Effects of BI9774 on metabolic endpoints in *Lep^ob^/Lep^ob^* AMLN mice

Treatment day 4 through 12 showed a significant decrease in absolute body weight between the high dose BI9774 group vs. the vehicle treated AMLN fed mice (Figure 3A). This is in line with reports of AMPK activation being associated with reduced body weight in mice on a high fat diet (Myers et al., 2017), (Cokorinos et al., 2017), (Pollard et al., 2019). No difference was seen in weight gain between groups regardless of the mice on the AMLN diet having a lower starting body weight than mice on LFD, (Figure 3B).

**Fig. 3:**
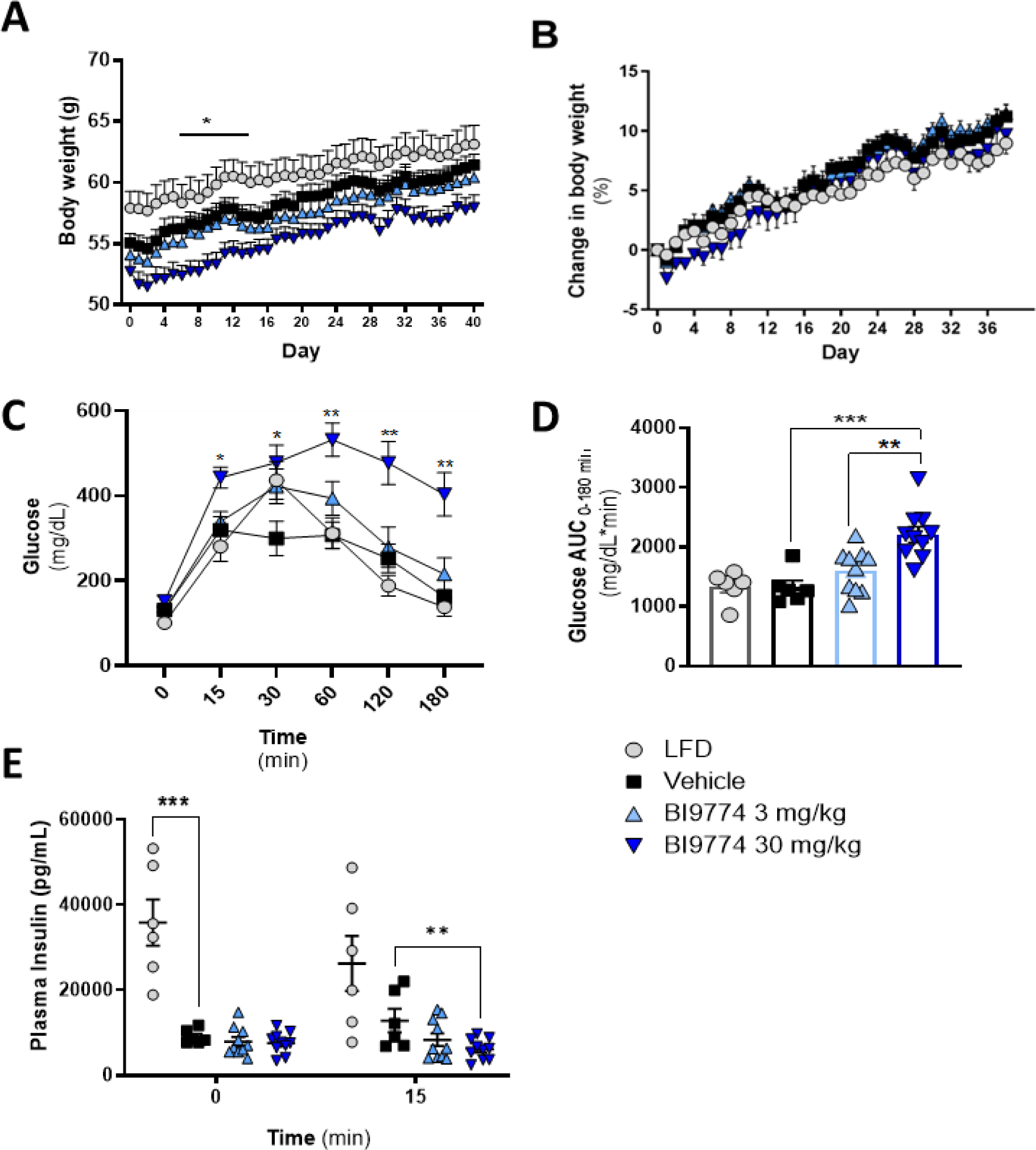
Effects of direct AMPK activation on body weight and glucose tolerance in *Lep^ob^/Lep^ob^* mice on AMLN diet. All groups consist of 9-week-old male *Lep^ob^/Lep^ob^* mice double housed and placed on either low fat diet (LFD) Research Diets #D09100304 n=8, or high trans-fat diet (AMLN) Research Diets #D09100301 for 14 weeks prior to dosing. AMLN fed groups consist of the following vehicle treated mice n=8, 3mg/kg treated mice n=10, and 30mg/kg treated mice n=10. **A)** Body weight measured daily. Analyzed by 2way ANOVA. **B)** Percent change in body weight compared to baseline showed no change in 2way ANOVA analysis. **C**) Blood glucose measured on a Breeze2 glucometer (Bayer, Pittsburgh, PA) following a 4h fast and 1.5g/kg intraperitoneal glucose bolus. Significance compared to vehicle treated mice **D**) Blood glucose area under the curve (AUC) analyzed with 1way ANOVA. **E)** Insulin measured using Meso Scale Discovery Mouse/Rat insulin kit (cat #K152BZC) at 0 and 15 minutes following a 1.5g/kg intraperitoneal glucose bolus. * p<0.05, ** p<0.01, ***p<0.001.

After 4 weeks of treatment (18 weeks on diet) an ipGTT showed impaired glucose tolerance in the 30 mg/kg dose compared to vehicle (Figure 3C-D). This was unexpected, but corelated with a dose-dependent reduction of plasma insulin at the 15 min timepoint (Figure 3E). An acute glucose tolerance test in diet-induced obese (DIO) C57Bl6J mice was performed to further investigate the unexpected findings on glucose tolerance and showed the expected dose-dependent improvement in glucose control. (Supplementary Figure 2).

### BI9774 improved liver metabolic endpoints in AMLN fed *Lep^ob^/Lep^ob^* mice

AMPK activation showed clear liver benefits in our NASH/NAFLD model. Terminal liver weight (expressed as % body weight) and liver fat (expressed as % of liver mass) were reduced 2.5% and 10% respectively in mice administered 30 mg/kg BI9774, which was equal or below that of the animals on LFD (Figure 4 A-C). Plasma ALT and AST levels both showed dose-dependent reduction trends by BI9774, although only ALT levels in the animals given 30 mg/kg were significantly below those of vehicle treated mice (Figure 4 D-E). We also measured expression of mRNA transcripts important in liver disease or regeneration. The mRNA transcript for CLEC2, *Clec2h*, showed a 15-fold increase in expression vs. vehicle after AMPK activation by 30 mg/kg BI9774. Expression of *Cyp46a1* and *Sult1c2* was lowered in livers of mice on the AMLN diet vs. LFD controls, and had a significant dose dependent increase with BI9774 treatment. There was no effect on lipid metabolism transcription co-factor *Ppargc1*a (Figure 4 F-G).

**Fig. 4:**
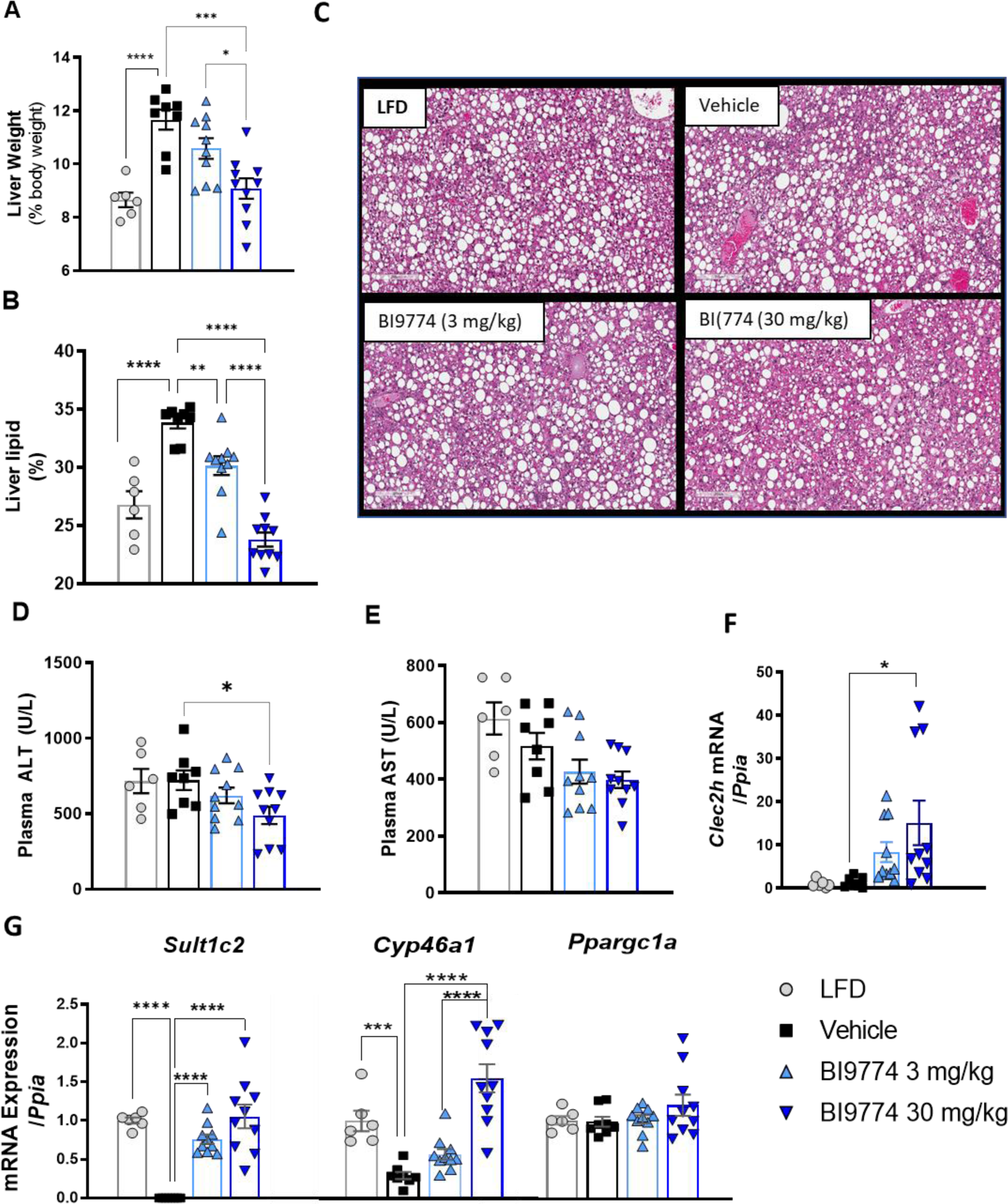
Effects of direct AMPK activation on liver biochemistry in *Lep^ob^/Lep^ob^* mice on AMLN diet: Terminal liver weight as a percentage of Body weight. **B)** Liver lipid was calculated as a % of total tissue mass. **C)** Histology of hemotoxlin and eosin-stained terminal liver. **D-E)** Terminal plasma AST and ALT collected via cardiac puncture on day 42 of treatment. **F-G)** Liver mRNA expression after 4 weeks of treatment with BI9774. Expression is normalized to *Ppia* and graphed relative to LFD control mice. Student’s t-test was used to compare low-fat diet vs. vehicle-treated controls. Drug effects vs. vehicle controls were determined using one-way ANOVA with Tukey’s post hoc test. * p<0.05 ** p<0.01, ***p<0.001, ****p<0.0001

### BI9774 reduced expression of fibrosis related mRNA, lowered circulating cytokines, and slowed progression of macrosteatosis

Collagen-1 immunopositivity increased in livers of mice on the AMLN diet compared to LFD controls while BI9774 administration did not alter Col1a1^+^ area, (Figure 5 A-B). Consistent with the Col1a1^+^ area data. We examined hepatic mRNA levels of collagen genes and *Timp1*. Groups treated with 30mg/kg BI9774 showed a 61% ± 4% decrease in expression of *Col1a1,* 59% ± 3% decrease in *Col1a2*, 55% ± 4% decrease in *Col4a1*, and a 55% ± 3% decrease in *Timp1* expression (Figure 5 G). *Col5a1* mRNA expression remained unchanged. Inflammation was quantified by IHC using pan macrophage marker CD68 (Chistiakov et al., 2017), (Figure 5 C) No significant changes in macrophage infiltration were observed, but a panel of proinflammatory cytokines showed significant reduction of IL-6 and IL-1β with BI9774 treatment. KC/GRO, an activator of proinflammatory and profibrotic genes (Stefanovic and Stefanovic, 2006), also showed a significant reduction with AMPK activation. (Figure 5 D-F) The 30 mg/kg dose slowed the progression of macrosteatosis, with only 30% of mice worsening vs. 50% of those given 3 mg/kg and 63% in the vehicle group. While the 30 mg/kg dose of BI9774 lowered the NAFLD activity score (NAS) from an initial 11.6 to 10.6, this was not significant. Other measures, biliary hyperplasia and lobular inflammation showed minimal or no changes although a trend for protection was seen (Figure 5 H-I) Circulating free fatty acids measured at the end of study also showed no significant change (Supplementary Figure 3A).

**Figure 5.**
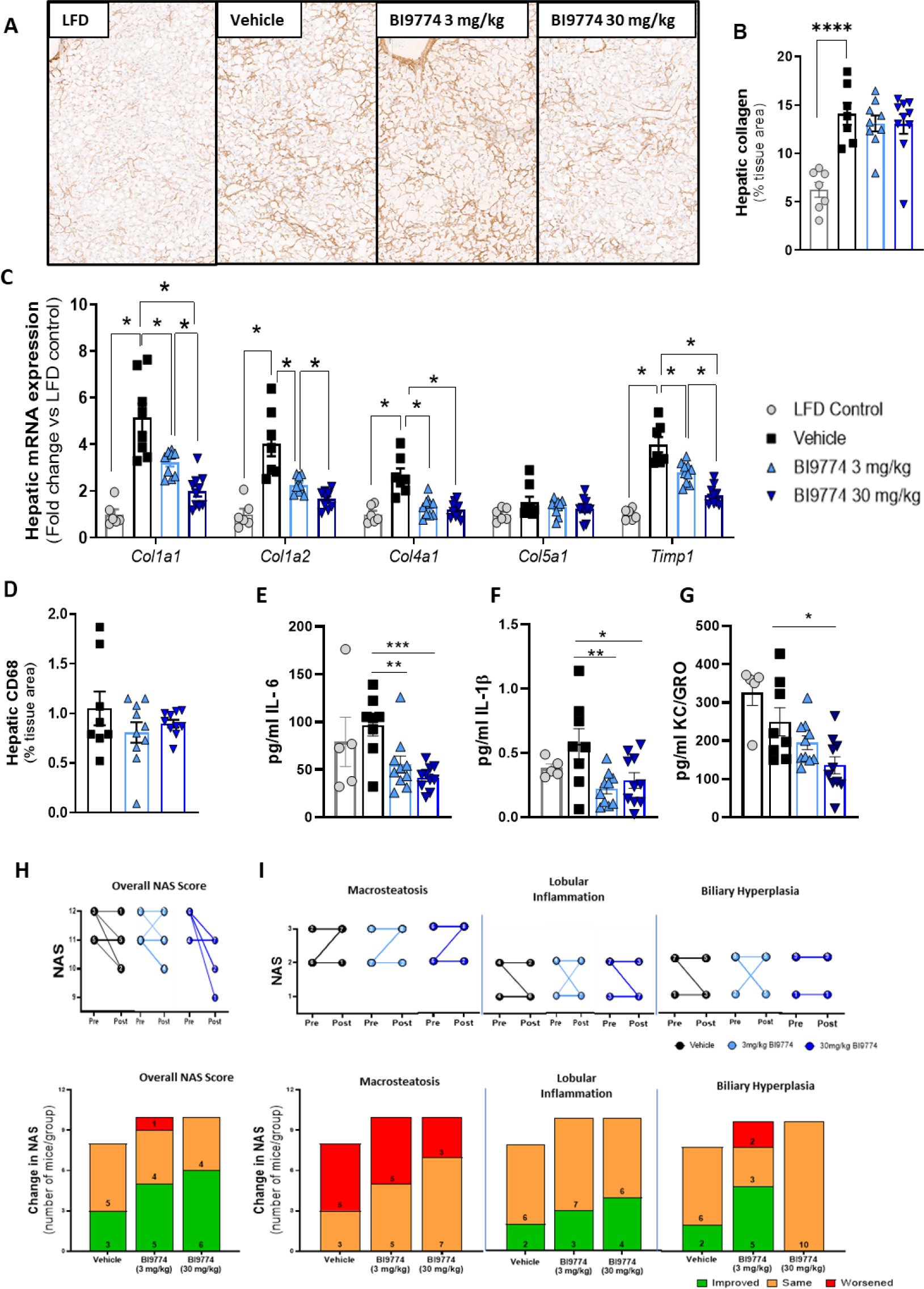
Effects of direct AMPK activation on liver histology in *Lep^ob^/Lep^ob^*mice on AMLN diet. **A&B)** Immunohistochemistry was performed using formalin-fixed, paraffin embedded liver sections. **C)** BI9774 treatment in *Lep^ob^/Lep^ob^* NASH mice was associated with significant reductions in the expression of fibrosis-related genes. Liver mRNA levels were measured by qPCR and are expressed as fold change relative to low-fat diet control mice. Student’s t-test was used to compare low-fat diet vs. vehicle-treated controls. Drug effects vs. vehicle controls were determined using one-way ANOVA with Tukey’s post hoc test (low-fat diet controls were not included). **D)** Inflammation was quantified using anti-CD68 (ab125212, Abcam)**-E-G)** Cytokines were measured using Meso Scale Diagnostics V-PLEX Proinflammatory Panel 1 Mouse Kit (Cat# K15048D) **H & I)** Overall NAS score and several markers of NASH were analyzed by Fisher’s Test. No significance was seen between pre and post dosing samples. * p<0.05 ** p<0.01, ***p<0.001, ****p<0.0001

### AMPK activation by BI9774 lead to cardiac hypertrophy and increased cardiac glycogen storage in AMLN fed Lep^ob^/Lep^ob^ Mice

We observed a 13.3% increase in heart weight relative to body weight, along with increased glycogen storage in mice given 30 mg/kg BI9774 (Figure 6A-D). *Col1a1* expression was affected dose dependently and significantly reduced in the high-dose BI9774 group vs. vehicle. The mice on AMLN diet showed reduced expression of *Nppa* and *Ankrd1* RNA compared to those on LFD, which was dose-dependently restored with BI9774 treatment (Figure 6E). Other genes found to play a role in hypertrophy including *Cyp46a1*, *Nfatc1*, *Ppargc1a*, and *Wdr1* showed no significant differences with treatment (Supplementary Figure 3B) (Santolini et al., 2018), (Molkentin, 2004), (Aihara et al., 2000), (Song et al., 2015), (Huang et al., 2019).

**Fig 6.**
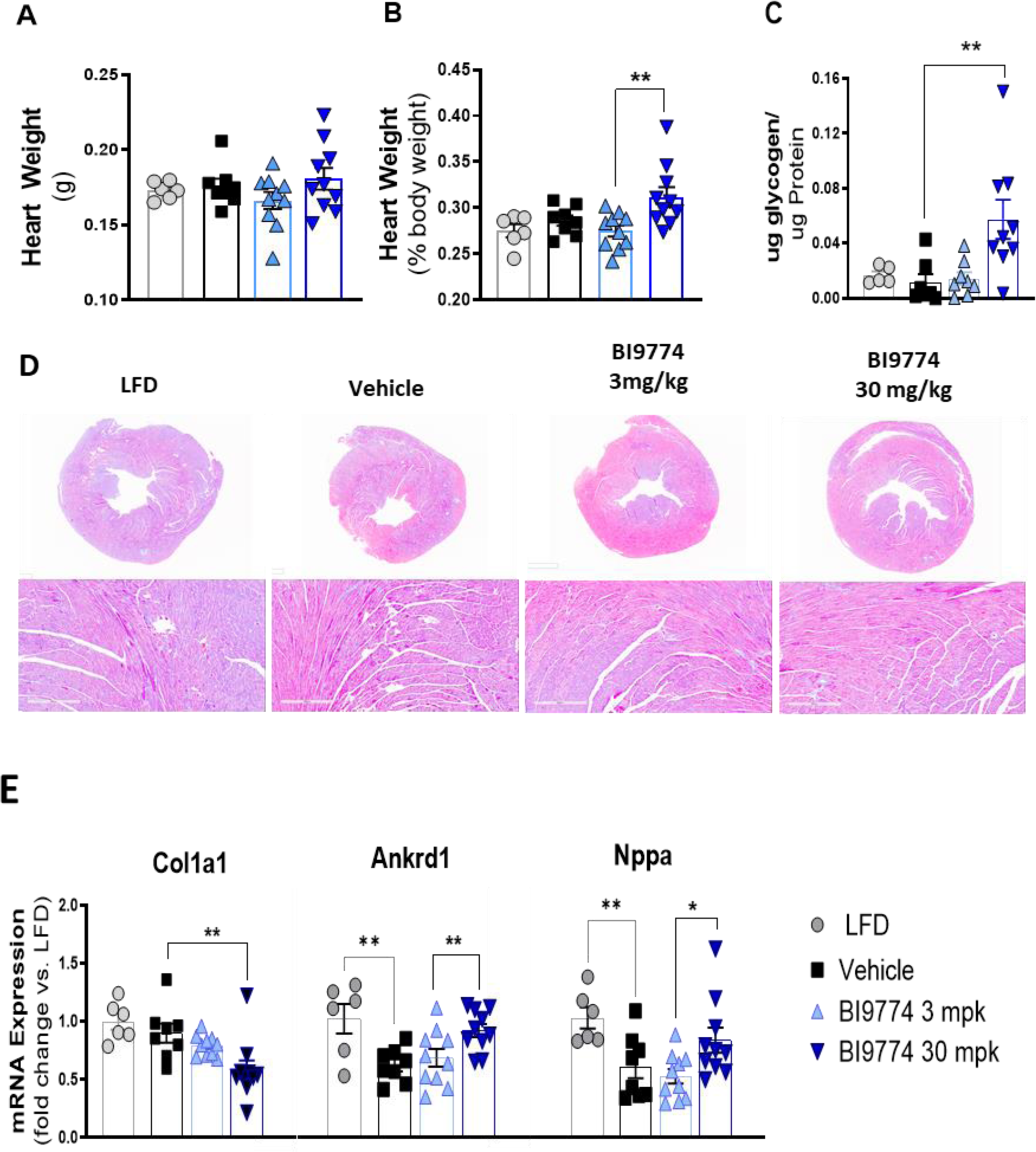
Effect of BI9774 on cardiac hypertrophy in *Lep^ob^/Lep^ob^* mice on AMLN diet: **A&B)** Six weeks of treatment with BI9774 showed a clear increase in heart weight to body weight ratio. **C)** There was a dose dependant increase in glycogen storage in the 30mg/kg group. **D)** Masson’s Trichome staining shows distinct thickening of the cardiac wall in mice treated with 30mg/kg BI9774 vs. the vehicle group. **E** Cardiac *Nppa* and *Ankrd1* expression was suppressed in mice on the AMLN diet. BI9774 restored expression in a dose dependent manner. Expression of *Col1a1* decreased below that of the LFD mice. Student’s t-test was used to compare low-fat diet vs. vehicle-treated controls. Drug effects vs. vehicle controls were determined using one-way ANOVA with Tukey’s post hoc test (low-fat diet controls were not included). *p<0.05, **p<0.01.

## Discussion

In silico and pharmacokinetic studies showed BI9774 to be a viable tool compound to assess AMPK activation as a target for NASH therapy *in-vivo*. A concentration response relationship between BI9774 and ACC phosphorylation in hepatocytes confirmed the cellular activity of the compound. We demonstrated that BI9774 engages AMPK at the relevant ADaM site, the key target site of other known AMPK activators, by generating a bound crystal structure. This was used as a proxy for target engagement. (Esquejo et al., 2018) (Myers et al., 2017) Pharmacokinetic studies in lean and *Lep^ob^/Lep^ob^*mice determined appropriate dosing to achieve sustained plasma exposure above the projected cellular EC_50_ for pACC levels in-vivo. This data was sufficient to conclude that BI9774 was a suitable tool compound for evaluating systemic AMPK activation’s effects on NASH.

Daily administration of BI9774 for 6 weeks at low (3 mg/kg) or high (30 mg/kg) doses in *Lep^ob^/Lep^ob^* mice with diet-induced NASH induced expected responses on several endpoints. The highest dose of BI9774 was associated with significant reductions in liver weight, hepatic steatosis, and plasma ALT levels in *Lep^ob^/Lep^ob^* mice with NASH and fibrosis. Low dose of BI9774 was associated with reduced liver weight and steatosis, indicating dose-dependent effects of AMPK activation on improved liver health. As food intake was not measured in this study it could be possible that reductions in caloric intake may have accounted for some of the body weight and/or liver axis improvements. It should be noted that others have reliably reported increased food intake in mice following AMPK activation (Andersson et al., 2004), and the lowest dose of BI9774 used here did not affect body weight significantly but did improve liver function. Therefore, we posit that any possible effects of AMPK activation in this model are likely due to direct activation of AMPK and not indirect effects resulting from changes in food intake.

The mechanisms/s whereby BI9774 improve liver lipid remain to be fully elucidated however several lines of evidence presented here suggest a key role for the regulation of cholesterol and lipid metabolism. Gene expression analysis of the liver revealed significant dose-dependent increases in mRNA expression of metabolic enzymes *Cyp46a1* and *Sult1c2. Cyp46a1* catalyzes cholesterol, steroid, and lipid metabolism (Lund et al., 1999), (Mast et al., 2003), (Dubaisi et al., 2018). *Sult1c2* mRNA is decreased in diabetic and obesity-induced steatosis experimental models (Almon et al., 2009), (Buque et al., 2010). These enzymes showed similar dose dependent restoration after treatment. Relative to the LFD control, Sult1c2 expression in the AMLN fed mice was restored from 0.11% in the vehicle treated group to 105% in the mice given 30 mg/kg BI9774. Cyp46a1expression fell to 0.3% in mice on the AMLN diet and increased to 155% with the high dosage. The lipid metabolism transcription co factor *Ppargc1*a, which plays a central role in the regulation of cellular energy metabolism (Liang and Ward, 2006), showed no effect from the diet or treatment and *Clec2h*, which is associated with improved liver steatosis by regulating Kupffer cell polarization (Wu et al., 2015), showed a strong dose dependent increase following treatment.

Surprisingly, a glucose tolerance test revealed worsened glucose excursion coupled with lower insulin levels in the mice treated with the high dose of BI9774. This contradicts previous studies on AMPK activators which reported improved or unchanged glucose appearance in the context of lower insulin levels (i.e., improved insulin action). (Carling, 2017) Upon further exploration using acute BI9774 treatment in C57BL6J diet induced obese (DIO) mice and observing the expected dose-dependent reduction in glucose area under the curve, we interpret the exaggerated glucose appearance in the *Lep^ob^/Lep^ob^* NASH mice with high BI9774 to be a unique function of pharmacological activation of AMPK in this model and diet rather than an accurate reflection of a worsened glycemic phenotype. While there is no definitive rodent model for metabolic/liver studies and there could be problems in interpretation, the *Lep^ob^/Lep^ob^* + AMLN diet model has been used consistently in preclinical studies for evaluating potential NAFLD and NASH therapies due to it exhibiting several hallmark features of NASH (steatosis, inflammation, and fibrosis) along with obesity and perturbed glycemic control (Trevaskis et al., 2012), (Hansen et al., 2017). The *Lep^ob^/Lep^ob^* mice studied here are aged compared to those typically used in metabolic studies. Combined with the NASH-inducing diet, the pancreatic changes that typically occur to combat the genetically induced hyperglycemia may be altered or lost. NASH mice exhibit lower fasting insulin compared to chow-diet control *Lep^ob^/Lep^ob^* mice and have no visible insulin secretion response upon glucose administration. Although, at less than 200 mg/dL, the fasting glucose levels observed here were relatively lower than previously seen in *Lep^ob^/Lep^ob^* mice (Sun, 2016). Further studies would be required to delineate this finding and understand its relevance to AMPK modulation.

Histologically, BI9774 was associated with subtle improvements in liver pathology driven largely by decreasing steatosis. This was reflected in the majority of animals exhibiting an improved NAS. Treatment significantly reduced the expression of multiple collagen genes. However, immunoreactive staining revealed no significant difference in collagen protein across AMLN groups, indicating temporal differences between AMPK-induced suppression of matrix production and resolution of the hepatic collagen. The effects on inflammation were also limited. We did see significant decreases in circulating in IL-6, IL-1β, and KC/GRO after BI9774 treatment but macrophage infiltration into the liver showed no changes between treated and control groups.

Our data also suggest that the beneficial metabolic and steatotic effects of BI9774 are at least somewhat separable from fibrosis regression. Although, it’s possible that dosing with BI9774 over a greater extent of time is required to reveal slowing or reversing of fibrosis. These effects are likely due to AMPK mediated inhibition of ACC, as hepatic suppression of ACC exerts clear benefits on liver metabolic phenotype, and inhibitors of ACC are being actively pursued as therapeutic therapies for NASH (Goedeke et al., 2018),(Kim et al., 2017). Recently Gluais-Dagorn et al reported that β1 specific AMPK activator PXL770 successfully reduced hepatic inflammation and fibrogenesis in C57Bl/6J *Lep^ob^/Lep^ob^* hyperphagic obese mouse model on a NASH diet mice after 8 weeks of treatment. It’s possible the extended treatment time or the subtype specific molecule is responsible for the decrease in inflammation. Changes in Col1a1^+^ area or existing fibrosis were not reported in their study (Gluais-Dagorn et al., 2022).

AMPK activation has shown evidence of adverse cardiac effects, such as being associated with cardiac hypertrophy in animal models and in humans with Wolff-Parkinson-White Syndrome (Myers et al., 2017), (Kim et al., 2014), (Yang et al., 2016). This syndrome, caused by a mutation in the AMPK γ2 subunit, leads to constitutive activation of AMPK. In accordance with previous observations, we also noted adverse pharmacological effects of BI9774 on cardiac tissue (Myers et al., 2017). Specifically, we observed a dose-dependent increase in glycogen storage, heart weight, and cardiac markers of hypertrophy. The latter two changes, which have been observed using several different AMPK activators and in different species, appear to be an on-target consequence of AMPK activation, at least via ADaM site activators (Santolini et al., 2018), (Aihara et al., 2000), (Lund et al., 1999). Recent reports suggest that extra-hepatic activation of AMPK is required for the full metabolic benefit of AMPK therapy (Myers et al., 2017). Whether restricting AMPK activity to the liver or fine-tuning potency to specific AMPK subunits can replicate the full therapeutic potential of AMPK activation in the context of NASH or alleviate the detrimental cardiac effects of systemic activation remains to be determined.

To conclude, 6 weeks of systemic AMPK activation by BI9774 in a *Lep^ob^/Lep^ob^*obese mouse model fed a NASH diet demonstrated significant improvements in liver biochemistry and suppressed hepatic collagen gene expression but did not improve histological fibrosis. Coupled with increased heart weight and glycogen storage, these data suggest that current systemic β1/ β2 AMPK activators may not be viable therapies for NASH-related fibrotic disease.

## Materials and Methods

### Caliper *In-Vitro* Assessment

AMPK (α1β1γ1 or α1β2γ1) phosphorylation of FAM SAMS peptide (5’-FAM-HMRSAMSGLHLVKRR-COOH – Perkin Elmer) was followed using a Caliper EZ reader (Perkin Elmer). AMPK (40nM), 2□M FAM SAMS peptide, and compound were preincubated for 5 mins at room temperature before addition of 10□M ATP(Sigma) in 100mM Hepes pH7.4 (Sigma), 5 mM MgCl_2_ (Sigma), 1mM DTT and 0.015% BRIJ-35(Sigma) to start the assay. Compounds were tested in a dose response range of 100□M to 100nM dissolved in DMSO. A final concentration of 1% DMSO was present in all assay wells. Plates were read after 60 minutes. Percent inhibition values were calculated using the equation (100 – ((sample response -background response) □ (control reading – background response) x 100). The percent inhibition values were negative for stimulators and were fitted to the logistic equation y = A+((B-A) □ (1+((C□x)^D))) where x = compound concentration (M) and y = percent Inhibition, A = bottom of curve, B = top of curve, C = midpoint of curve (EC_50_), D = slope. pEC_50_ = -log_10_(EC_50_) and represents the concentration of compound giving half the maximum stimulation.

### BI9774 Selectivity

Selectivity studies were contracted to Thermo Fisher to run their panel screen of 408 protein kinases which can be found at https://www.thermofisher.com/uk/en/home/industrial/pharma-biopharma/drug-discovery-development/target-and-lead-identification-and-validation/kinasebiology/kinase-activity-assays.html against 1 μM of BI9774 (Thermo Fisher Scientific, UK); and in dose response against a panel contracted out to Eurofins consisting of 70 other custom selected diverse targets from their catalogue https://www.eurofinsdiscoveryservices.com/services/adme-tox/in-vitro-toxicology/ (Eurofins Cerep, Poitiers, France)

### AMPK Protein purification

DNA sequences for full-length human AMPK subunits were optimized for *E. coli* codon usage; and cloned into tri-cistronic pET-based plasmids, with a non-cleavable hexahistidine tag fused to the N-terminus of the alpha subunit. Protein was expressed in *E. coli* strain BL21 (DE3); and purified by Ni^2+^-IMAC followed by size-exclusion chromatography. Purified protein was incubated with CaMKK1 (molar ratio of 1:20) and 0.5 mM ATP in 25 mM Tris-HCl pH 8.0, 300 mM NaCl, 1 mM TCEP, 10 % Glycerol, 1 mM CaCl_2_, and 10 mM MgCl_2_ for 16 h at 18 C. CAMKK was removed by Ni^2+^-IMAC. Stoichiometric phosphorylation of the beta subunit and was confirmed by mass spectrometry, whereas the phosphorylation of the alpha subunit was not detectable. Final protein buffer was 50 mM Tris-HCl pH 8.0, 300 mM NaCl, 1 mM TCEP. Protein intended for enzymology was further supplemented with 10 % glycerol. Human CaMKK1β was expressed in E. coli as an N-terminal GST-fusion protein; and purified by glutathione-sepharose affinity chromatography followed by size-exclusion chromatography.

### X-ray Crystallography

AMPK α2β1γ1 protein was stored at -80 °C in aliquots at a concentration of 5 mg/ml (38 µM) and in a buffer containing 50 mM Tris pH 8.0, 300 mM NaCl, 1 mM TCEP. For crystallization, a protein aliquot was thawed and mixed with a 4-fold molar excess of AMP (100 mM stock in water), a 1.2-fold molar excess of staurosporine (10 mM stock in DMSO) and a 3-fold molar excess of the activator (50 mM stock in DMSO). After 30-minute incubation on ice, insoluble material was removed by centrifugation. The pre-formed complex was crystallized using the hanging drop vapor diffusion method at 20 °C by mixing 2 µl of the protein solution with 1 µl of the reservoir solution containing 8 % PEG3350, 0.3 M guanidine hydrochloride, 0.1 M PIPES buffer pH 7.2. Polyhedral crystals formed within 3—7 days; they were cryo-protected by 1-second immersion in reservoir solution supplemented with 20 % ethylene glycol, and frozen by rapid immersion in liquid nitrogen. Crystals belonged to space group *P*2 (McCoy et al., 2007) and contained 2 AMPK heterotrimers per asymmetric unit. X-ray diffraction experiments were conducted at Diamond Light Source beamline I02. Diffraction images were integrated with DIALS (Winter et al., 2018) and scaled with AIMLESS (Evans and Murshudov, 2013) the structure was solved by molecular replacement with PHASER (McCoy et al., 2007) using one AMPK trimer from PDBID 4CFE as the search model. The initial solution was refined by manual model building in Coot (Emsley et al., 2010) and automatic refinement using Buster (Bricogne G). Restraints for staurosporine and the AMPK activator were generated using GRADE (Smart OS, 2011). Data collection and refinement statistics are summarized in Supplementary table 1.

### Cell-based pACC AssayCell-based pACC Assay

HTRF assay kit 64ACCPEG from CisBio was used to quantify endogenous levels of phospho-ACC-Ser79 according to manufacturer’s instructions with the following modifications: HepG2 cells were grown in EMEM (without phenol red) + 10% FCS, 1% glutamine, 1% NEAA, and 1% sodium pyruvate. Small volume white 384-well plates (Greiner 784075) were pre-loaded with test compounds. HepG2 cells (8000 in 5 µl growth media) were plated per well and plates were incubated for 1 h at 37⁰C, 5% CO2. Supplemented lysis buffer from assay kit 64ACCPEG (2 µl) was added per well and plates were incubated for a further 30 min at room temperature. 1 µl of the mixture of the 2 HTRF conjugates from assay kit 64ACCPEG was added and plates were incubated overnight at room temperature. The HTRF conjugates are each diluted 1/20 in detection buffer and pre-mixed just prior to use. The next day a Pherastar plate reader was used to read fluorescence emission at 665 nm and 620 nm.

### In vivo studies

Animal studies were approved by the Institutional Animal Care and Use Committee at AstraZeneca (Gaithersburg, MD) in accordance with the Guide for the Care and Use of Laboratory Animals. Animals were housed in standard caging at 22°C on a 12h light: dark cycle. Animal studies are reported in compliance with the ARRIVE guidelines. (Percie du Sert et al., 2020)

### Pharmacokinetic studies

Pharmacokinetic (PK) studies were conducted in male *Lep^ob^/Lep^ob^*mice n=3 per group. BI9774 was administered orally or intravenously as indicated as a solution containing 5% DMSO, 95% sulfobutylether β-cyclodextrin (SBE-ß-CD) (30% w/v) in water. Blood samples obtained postdosing were collected in plastic micro centrifuge tubes containing EDTA-K2 and centrifuged at 4000 × g for 5 min to obtain plasma. The plasma BI9774 concentration was determined using liquid chromatography-tandem mass spectrometry (LC-MS/MS) and monitoring transition from m/z 541 to 383 in ESI-positive mode. PK parameters were calculated by non-compartmental model using Phoenix WinNonlin v. 6.1 (Pharsight Corp, Certara, St. Louis, MO, USA).

### Glucose tolerance test after acute treatment with BI9774 in DIO mice

C57Bl/6 diet induced obese (DIO) pre-conditioned male mice arrived from Charles River Laboratories at 12 weeks of age and were singly housed on 60% kcal high-fat diet (D12492 Research Diets, New Brunswick, NJ) and automatic water. Animals were on diet for 1.5 months before arriving and stayed on the diet inhouse for another 2 months until they reached optimal weight. The morning of the study, mice were sorted by body weight into groups of 8 and fasted for 4 hours. Then baseline glucose measurements were taken (Breeze2 glucometer, Bayer, Pittsburgh, PA) and BI9774 (1, 3, 10, or 30mg/kg), vehicle, or 75 mg/kg metformin (Sigma) was administered by oral gavage in a solution containing 0.25% HPMC, 5% Tween80, 5% DMSO. One hour later blood glucose measurements were again taken followed by 1.5 g/kg glucose bolus, also administered by gavage. Glucose measurements followed at 15, 30, 60, and 120 minutes. All measurments were taken via tail bleed. Analysis was performed as glucose area under the curve (AUC) comparisons to control treated mice using one-way ANOVA test with Dunnett post-hoc test. Calculations using Graph Pad Prism 7 (GraphPad Software, San Diego, CA)

### Assessment of BI9774 on NASH Endpoints in *Lep^ob^/Lep^ob^*Diet-Induced NASH Model

Male *Lep^ob^/Lep^ob^* mice (Jackson Laboratory, Bar Harbor, ME), aged ∼8 weeks were placed on either a high *trans*-fat, high fructose, and 2% cholesterol Amylin Liver NASH Diet (AMLN) D09100301, or a nutrient matched diet with no premix, no additional fructose, and no added cholesterol other than that supplied by lard (LFD) D09100304., (Research Diets, New Brunswick, NJ) for 14 weeks. Approximately 2 weeks prior to study start, all mice on AMLN diet received a baseline liver biopsy under isoflourane anaesthetic, as previously described (Oldham et al., 2019). Mice were allowed to recover and on day -3 (first dose was performed on day 0). Mice on AMLN diet had a non-fasted blood sample collected via retroorbital bleed and then randomized to drug treatment groups based on fibrosis stage, body weight, blood glucose (Breeze2 glucometer, Bayer, Pittsburgh, PA), and plasma ALT (Cobas c-111, Roche Diagnostics USA, Indianapolis, IN). Mice on LFD were administered vehicle throughout the study. Mice on AMLN diet were administered either vehicle (0.25% HPMC, 5% Tween80, 5% DMSO), or BI9774 at doses of 3 mg/kg or 30 mg/kg, A sample size calculation was undertaken prior to conducting this study, resulting in a biostatistician recommendation of n=10 for the treatment groups to show a significant effect on metabolic endpoints with 80% power. In addition, based on well-established baseline results observed from our historical in-house studies, we were able to reduce the sample size of both vehicle-treated control groups. In our labs, the variability of the NASH measurement from the AMLN+Vehicle group was consistently smaller compared to drug-treated groups. Further, we have observed a large NASH effect between AMLN + Vehicle and LFD + Vehicle, permitting a smaller LFD sample size (n=6) for this study in accordance with 3R guidelines.

Vehicle and BI9774 were administered via oral gavage (5 mL/kg) once daily just prior to lights off for 6 weeks. Body weight was measured daily. An intraperitoneal glucose tolerance test using a 1.5 mg/kg glucose bolus was performed on day 28 after a 4 h fast measurements were recorded at 15, 30, 60, and 120 minutes. On day 42 mice were euthanized via CO2 inhalation in the non-fasted state, blood was collected via cardiac puncture; livers and hearts were excised, weighed, and processed for further analysis.

### Histological Analysis of Liver and Heart Tissue

Livers and hearts were fixed in 10% neutral buffered formalin for 24 h. Paraffin-embedded tissue sections were stained with hemotoxylin and eosin (liver) or Masson’s trichrome (heart) using standard procedures. Histological assessments of the liver were conducted by a pathologist under blinded conditions. A modified scoring system, based on the Brunt and Kleiner NAFLD activity score (NAS), previously developed and validated to enable a more reproducible and semi-quantitative assessment of murine liver, was used to quantify various parameters of the liver phenotype. (Jouihan et al., 2017) Immunohistochemistry was performed using a Ventana Discovery ULTRA Staining Module (Ventana Medical Systems, Tucson, AZ). Formalin-fixed, paraffin embedded liver sections were stained with anti-collagen type1 A1 (1310-01 Southern Biotech, Birmingham, AL). Inflammation was quantified using anti-CD68 (ab125212, Abcam)

### Measurement of Plasma Biochemicals

Terminal blood was collected in EDTA-coated tubes and centrifuged at 10,000 x g for 10 min. Plasma was collected and analyzed for triglycerides, total cholesterol, alanine aminotransferase ALT and aspartate aminotransferase (AST) levels using a biochemical analyzer (Cobas c-111, Roche Diagnostics, Indianapolis, IN). β-Hydroxybutyrate was measured using the β-Hydroxybutyrate (β-HB) Assay kit (C) (BioVision Milpitas, California). Cytokines were measured using Meso Scale Diagnostics V-PLEX Proinflammatory Panel 1 Mouse Kit Cat# K15048D Rockville, Maryland.

### Liver Lipid Quantification

Total hepatic lipids were measured in liver samples using a Bruker LF-90II minispec system (Bruker Biospin Corporation, Billerica, MA). The data are expressed as the percent lipid relative to the total tissue mass.

### RNA Isolation and Real-Time PCR

Total liver RNA was isolated using Qiagen RNeasy® columns (Qiagen, USA) according to the manufacturer’s protocol, including on-column DNA digestion using DNase I. Equal amounts of RNA were reverse transcribed to cDNA using SuperScript III First Strand cDNA synthesis kit (Invitrogen, Carlsbad, CA). Real-Time PCR was performed on a QuantStudio-7 Flex System (Applied Biosystems, Foster City, CA) using Applied Biosystems TaqMan Fast Universal PCR Master Mix and TaqMan probes (specific assays available on request). Each sample was assayed in triplicate and quantified using the 2-ΔΔCT method normalized to endogenous control *Ppia*. Data are expressed as fold change relative to low-fat diet control mice.

### Materials

The synthesis of BI-9774 was performed by Pharmaron Beijing Co. (Beijing, P.R. China) and is described in the following International patent application: WO2015007669A1, Example 2 (Himmelsbach F, 2015). Anti-CD68 was purchased from Abcam (ab125212). Anti-collagen type1 A1 from Southern Biotech (1310-01) was used for Collagen quantification. Pharmacological agents were purchased from Sigma-Aldrich unless stated otherwise. Lep^ob^/Lep^ob^ mice were purchased from Jackson Laboratories (Bar Harbor, ME).

### Data and Statistical Analysis

Fisher’s test analysis was run in R 4.0.0 (R_Core_Team, 2020) All other statistical analyses were carried out using GraphPad Prism 7 (GraphPad Software, San Diego, CA). The data were analyzed via either Student’s t-test, one-way or two-way ANOVA with appropriate post hoc tests as indicated in the figure legends. Values of p<0.05 were considered to represent statistically significant differences. The data and statistical analysis comply with the recommendations on experimental design and analysis in pharmacology (Curtis et al., 2018).

## Acknowledgements

We greatly appreciate the expertise of Steven Novick and Jean-Martin Lapointe and thank them for their contribution to this publication. We also thank Xiaohui Pei, Haijun Wang, and Guang He at Pharmaron for their support in the synthesis of BI9774.

## Conflict of interest Declarations

All authors are or were at the time of this work employees of AstraZeneca LP or AstraZeneca PLC. Authors also declare stocks in AstraZeneca

## Funding Statement

All research was funded by AstraZeneca LP

## Ethics Statement

Animal studies were approved by the Institutional Animal Care and Use Committee at AstraZeneca (Gaithersburg, MD)

## Data Availability Statement

The data that support the findings of this study are available from the corresponding author upon reasonable request. Some data may not be made available because of privacy or ethical restrictions.

## Supporting Information

**S1 Table.**
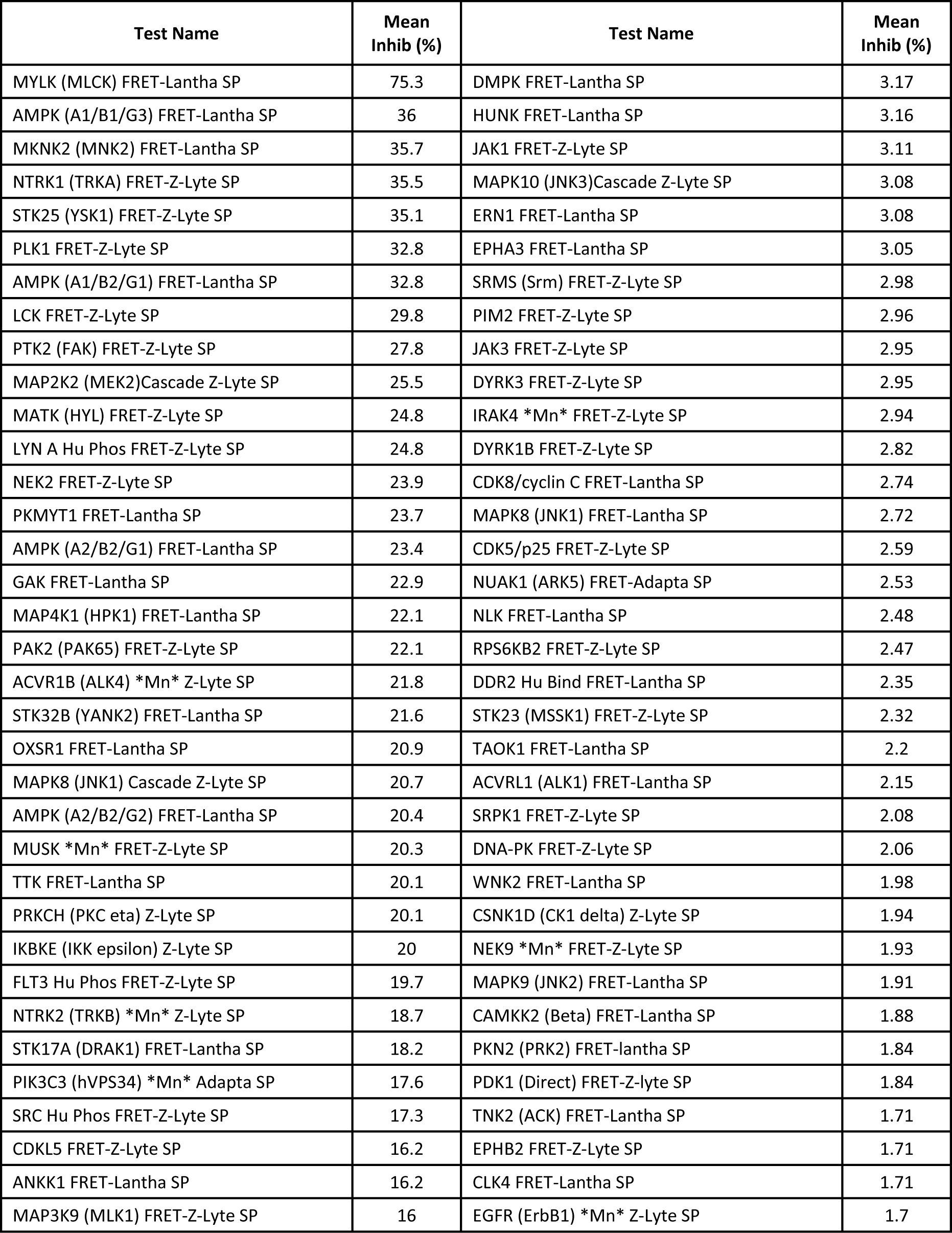
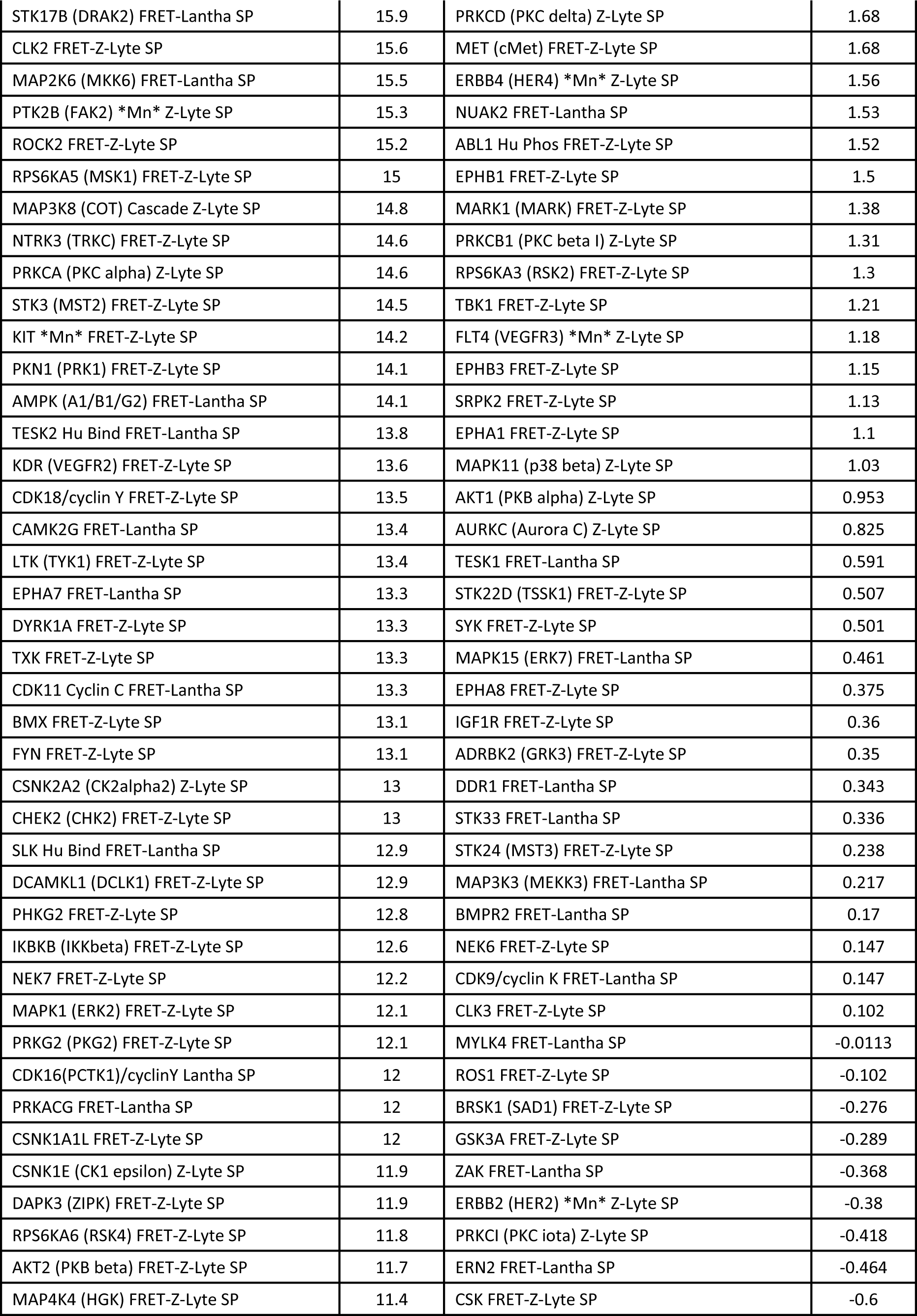
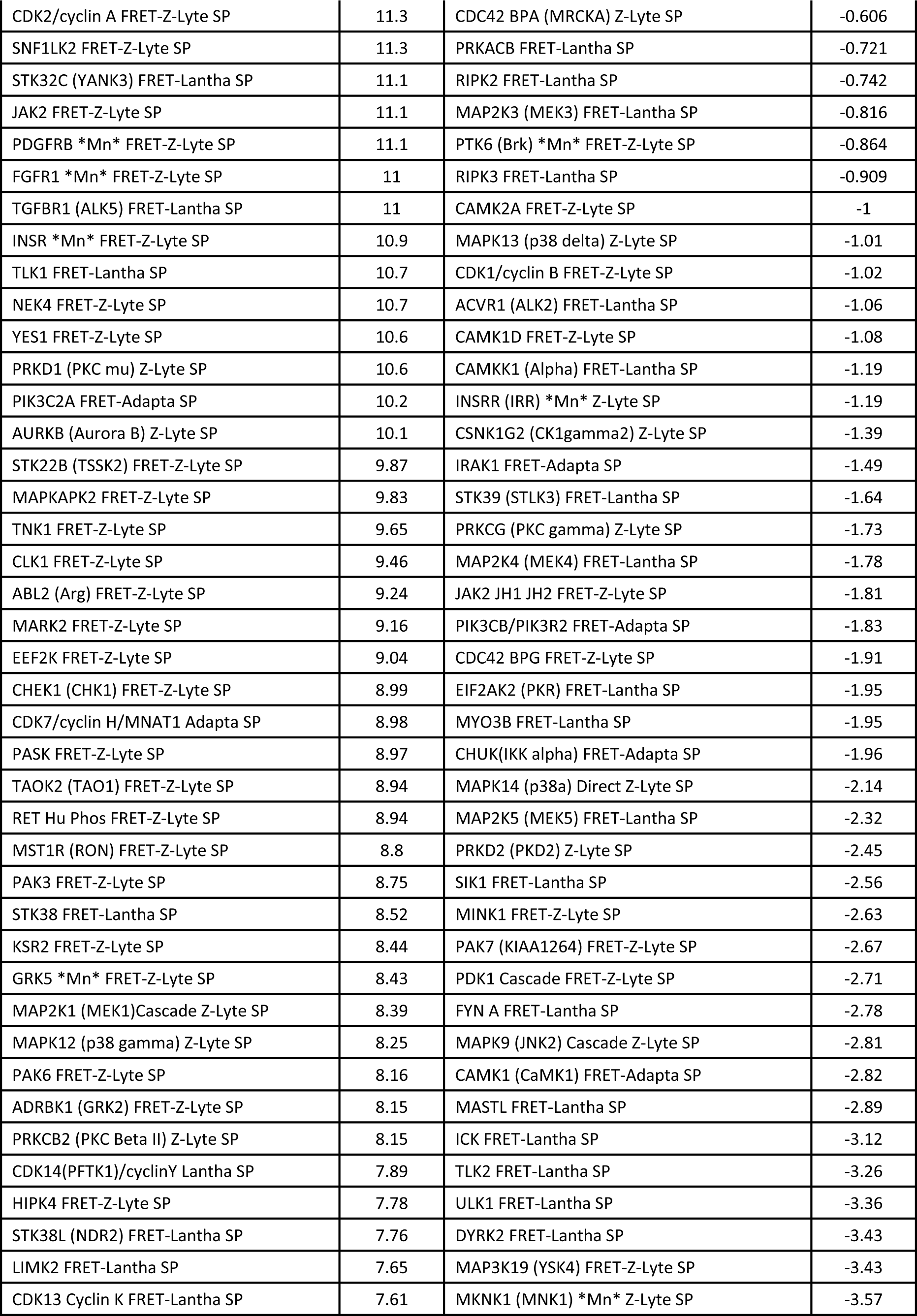
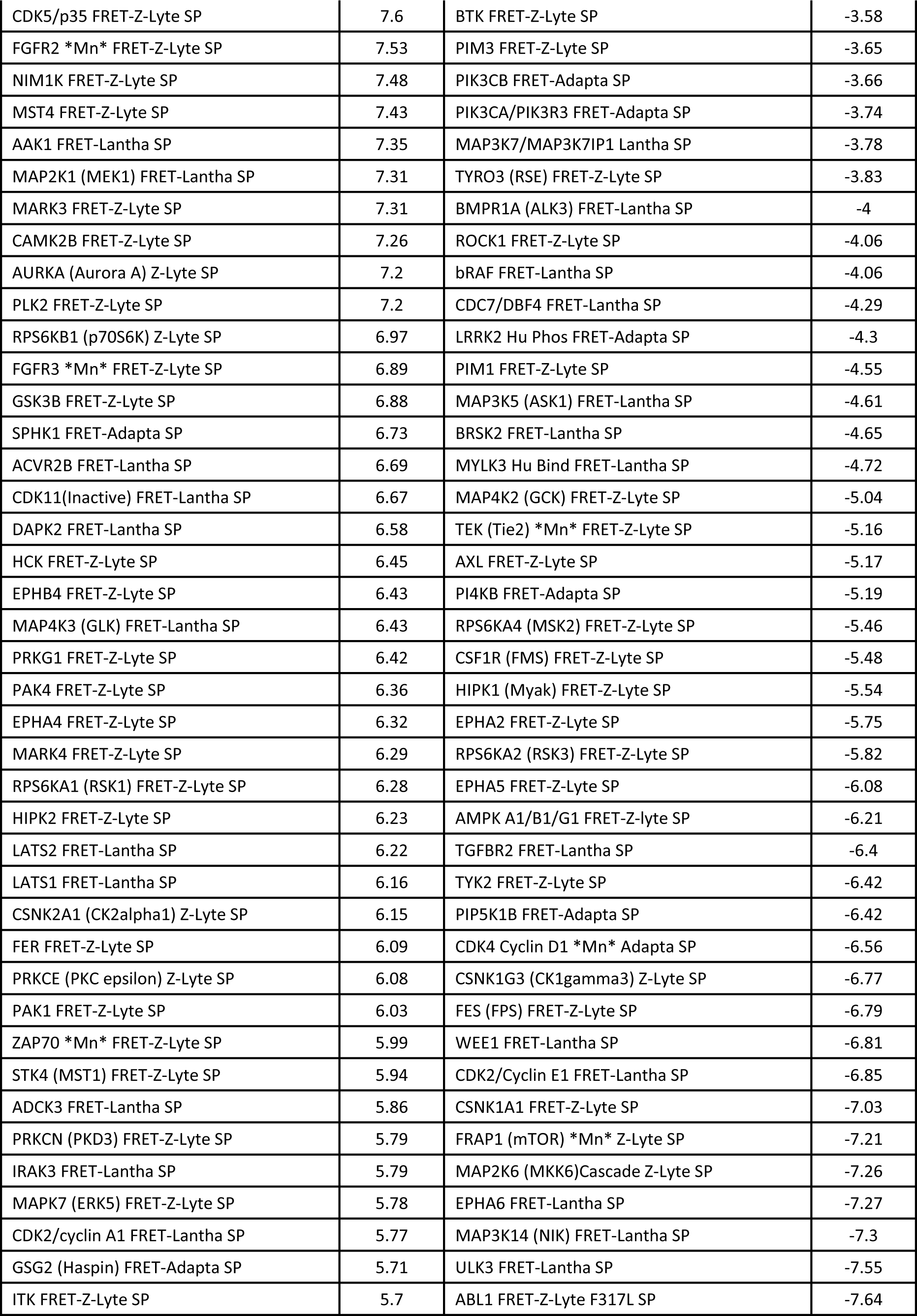
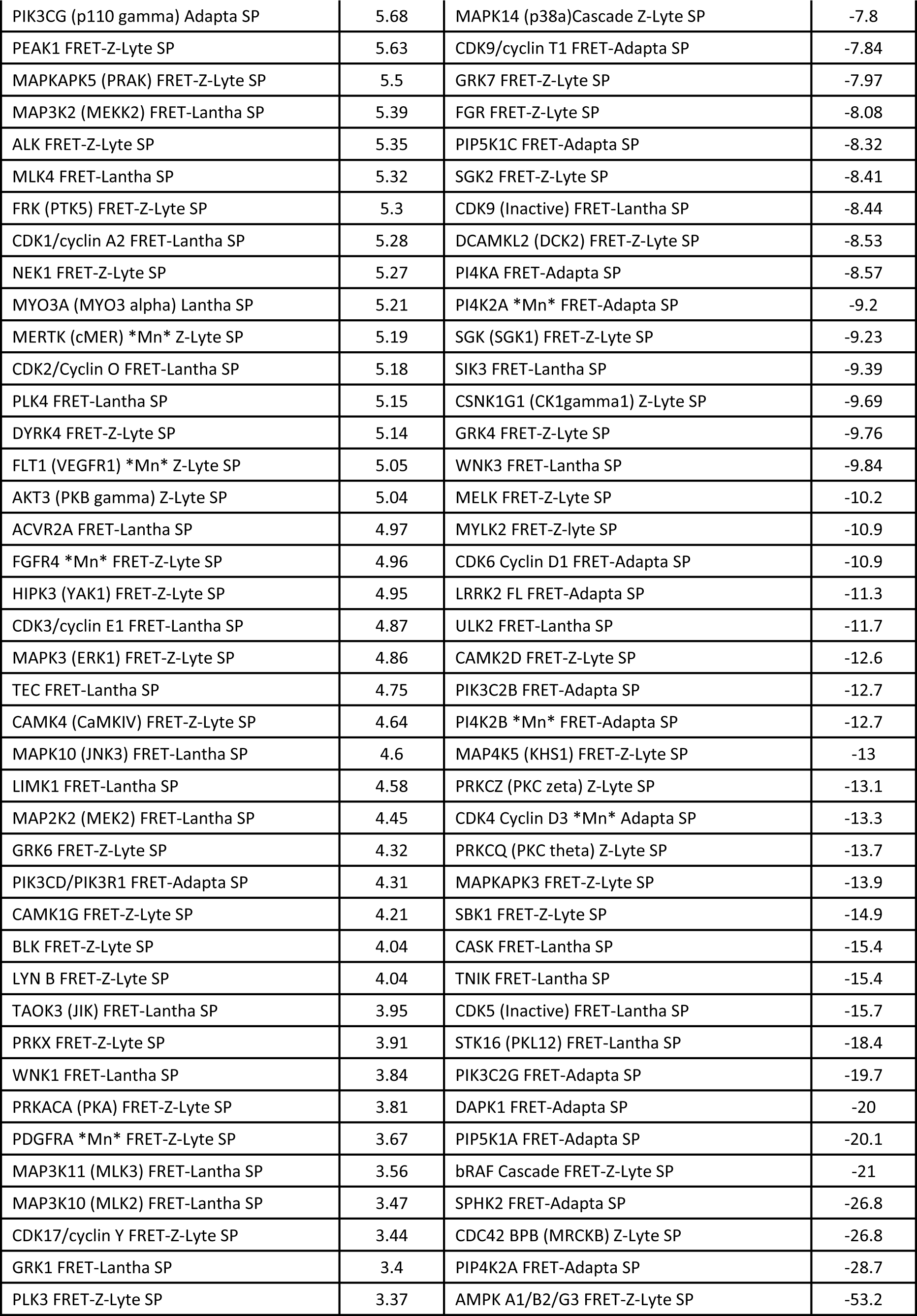
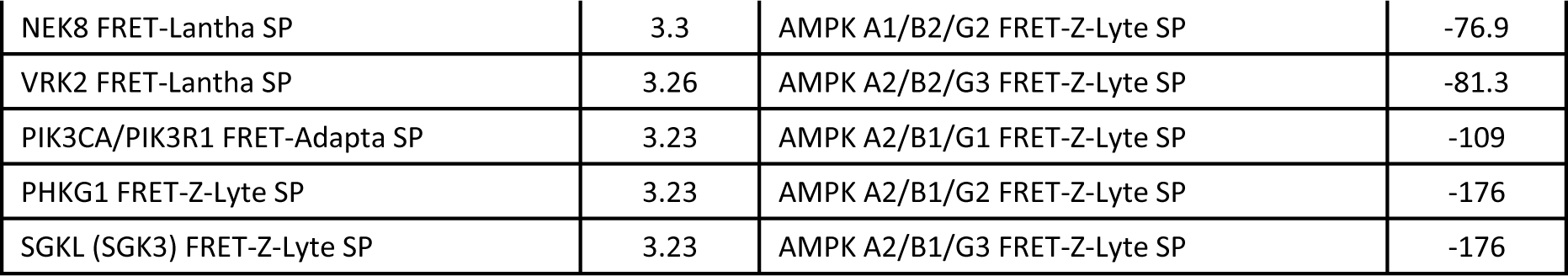
Results from Thermo Fisher’s protein kinase selectivity panel against 1uM BI9774.

**S2 Table.** X-ray crystallographic data collection and refinement statistics.

| AMPK $\alpha 2\beta 1\gamma 1$ with<br>staurosporine, AMP and<br>activator BI9774 | |
| --- | --- |
| PDB ID | 8BIK |
| **Data collection** |  |
| Space group | $P2_1$ |
| Cell dimensions |  |
| $a, b, c$ (Å) | 75.31<br>127.97<br>139.22 |
| $\alpha, \beta, \gamma$ (°) | 90.0<br>92.91<br>90.0 |
| Resolution (Å) | 75.22-2.50 (2.56-2.50) |
| $I / \sigma I$ | 8.9 (1.4) |
| CC 1/2 | 0.988 (0.546) |
| Completeness (%) | 99.9 (99.9) |
| Redundancy | 3.3 (3.4) |
| **Refinement** |  |
| Resolution (Å) | 75.22-2.50 |
| No. reflections | 300129 |
| $R_{\text{work}} / R_{\text{free}}$ | 0.181 / 0.215 |
| No. atoms |  |
| Protein | 15038 |
| Staurosporine | 70 |
| Activator 455 | 74 |
| AMP | 115 |
| Water | 482 |
| **B-factors** |  |
| Protein | 67.0 |
| Staurosporine | 41.9 |
| Activator 455 | 57.7 |
| AMP | 68.9 |
| Water | 58.0 |
| **R.m.s. deviations** |  |
| Bond lengths (Å) | 0.01 |
| Bond angles (°) | 1.13 |

**Supplementary Figure 1.**
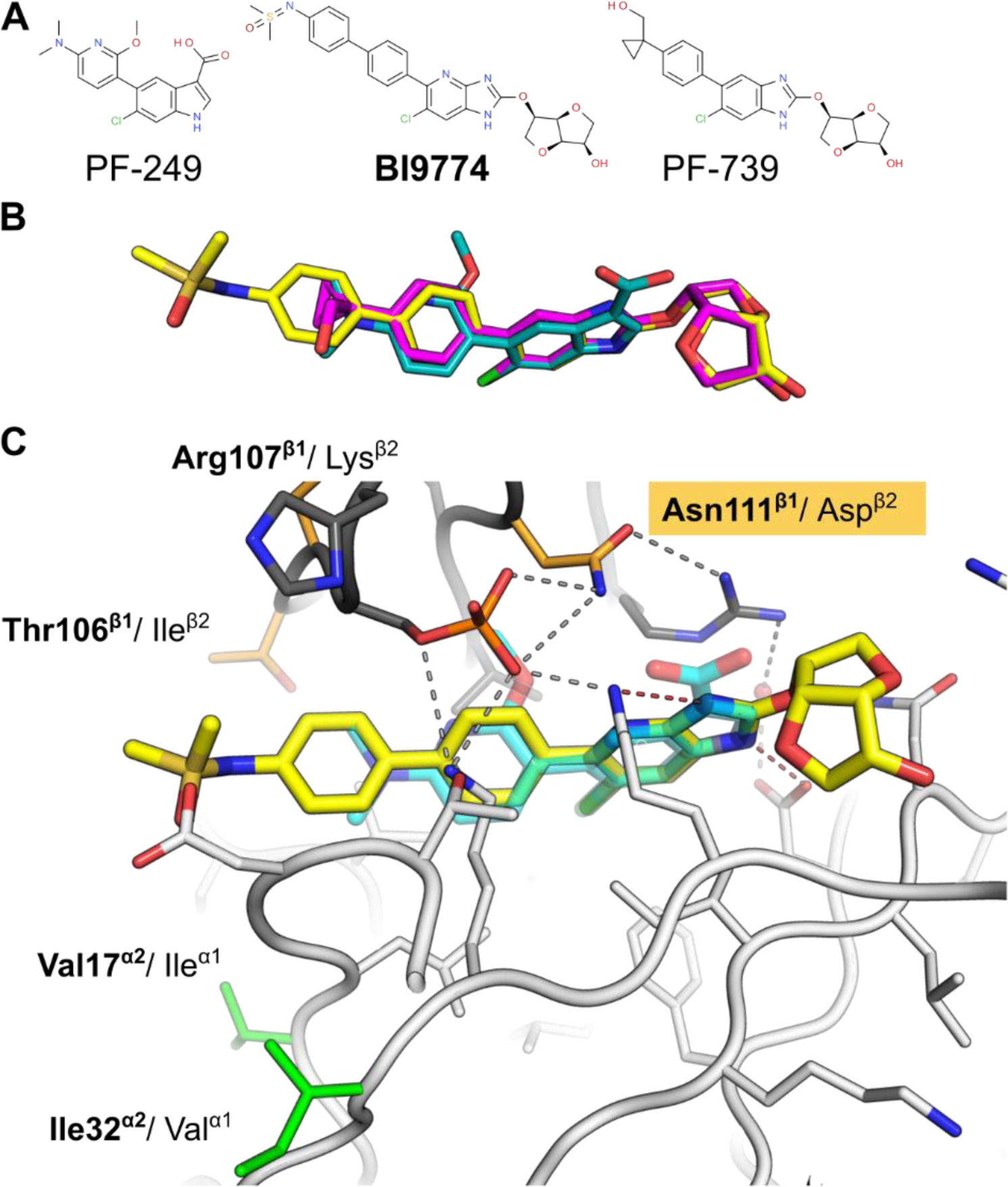
Activator BI9774 avoids the isoform-specific agonism caused by isoform sequence differences in the allosteric drug and metabolite (ADaM) site. **A)** Chemical structure of β1-selective AMPK activator PF-249, and pan-specific agonists BI9774 and PF-739. **B)**. Similarity in the binding pose of AMPK activators: BI9774 (yellow, PDB: 7A07), PF-739 (cyan, PDB: 5UFU) and PF-249 (magenta, PDB: 5T5T). **C)**. Crystal structure of V455 with AMPK α2β1γ1; PF-739 was super-imposed from PDB: 5T5T. Isoform sequence differences in the alpha and beta subunits are highlighted as coloured sticks (orange for beta, and green for alpha). Within direct interaction distance to the activator, the only difference is amino acid 111 in the beta subunit: asparagine in β1 and aspartic acid in β2. The reduced affinity of PF-249 for β2 is attributed to electrostatic repulsion between the negative charge on Asp111^β2^ and the acid moiety on PF-249. The uncharged group of 455 conveys comparable affinity for both beta isoforms.

**Supplementary Figure 2:**
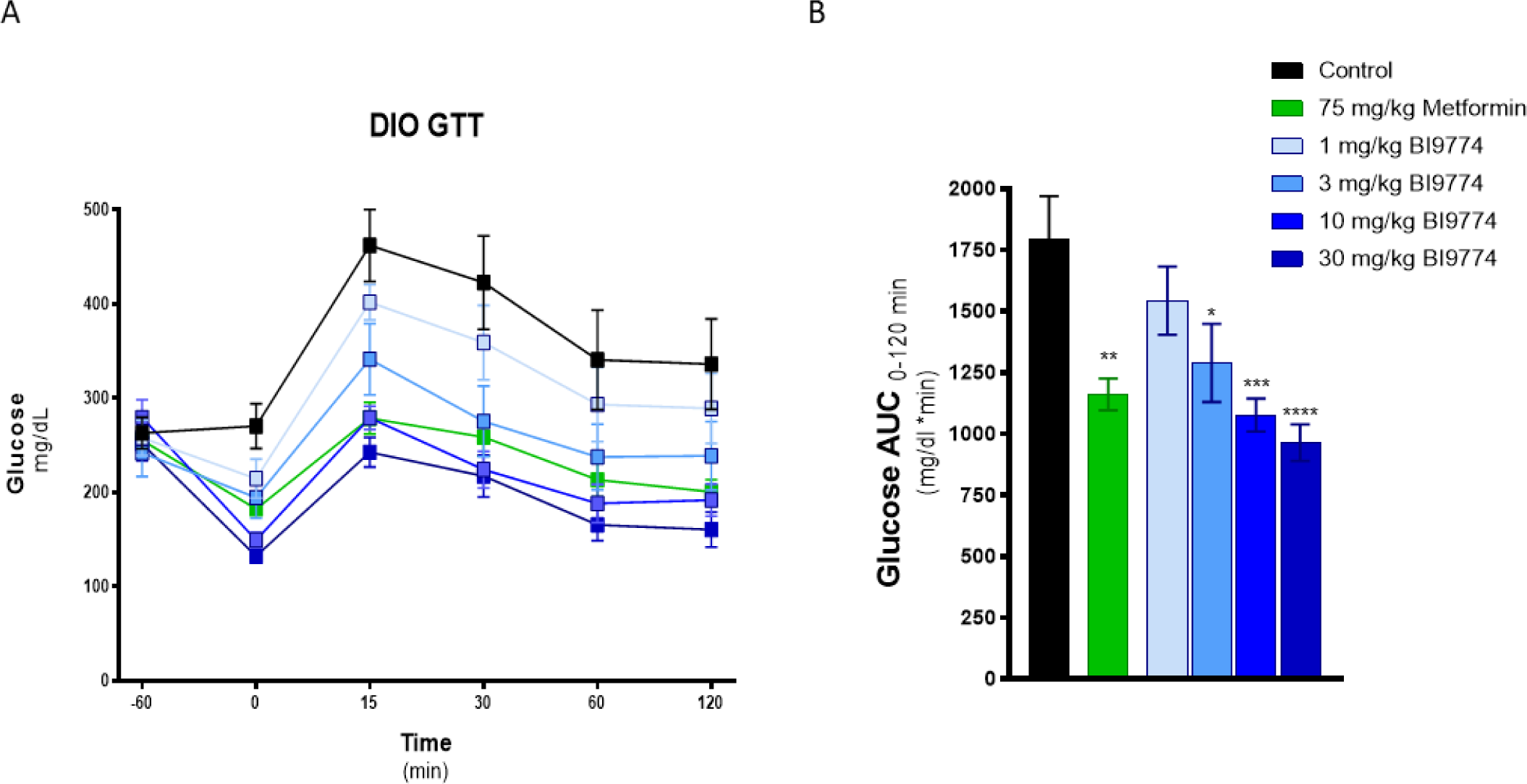
Acute BI9744 increases glucose tolerance dose dependantly in DIO mice. **A)** Blood glucose concentrations of DIO mice at each of the timepoints indicated. Comparisons are vs. Control mice calculated by two-way ANOVA with Tukey’s repeated measures post hoc test, **B)** Calculated glucose area under the curve (AUC). Analysis was performed as comparisons to control treated mice using one-way ANOVA test with Dunnett post-hoc test. * p< 0.05, ** p<.01, *** p<.001, **** p<.0001.

**Supplementary Figure 3:**
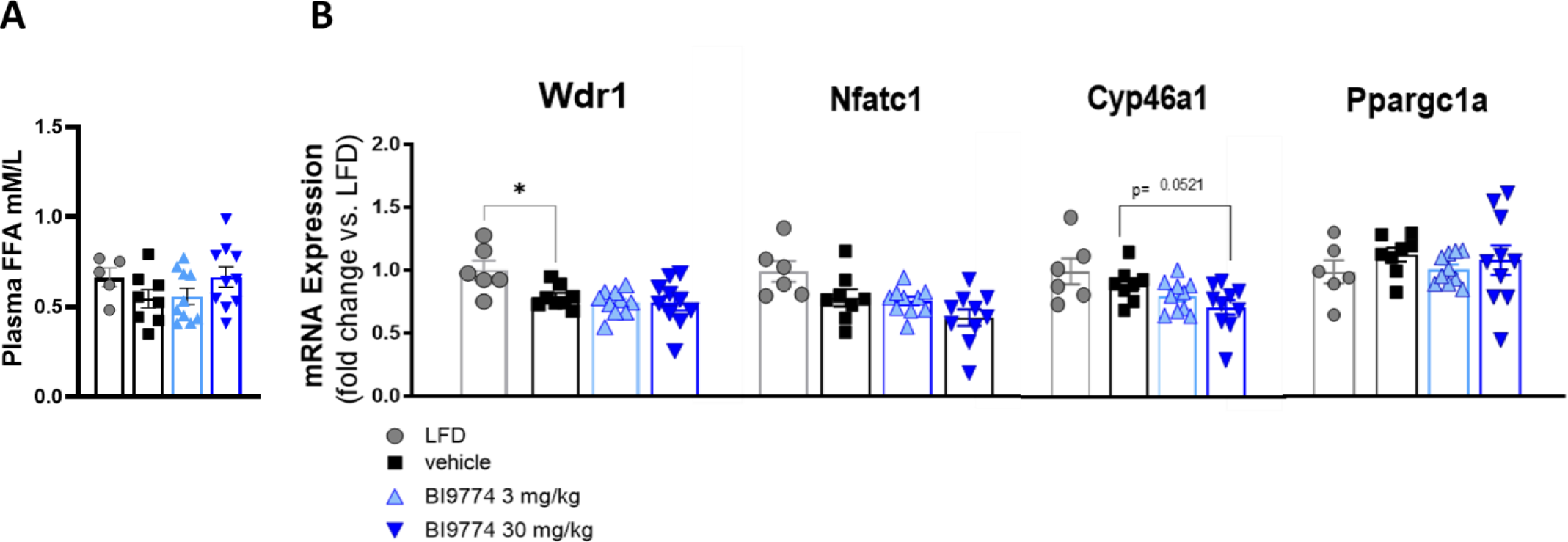
Some cardiac expression of hypertrophic markers remained unchanged. **A)** Plasma FFA were quantified using Wako FNEFA-HR Assay (VWR cat# 99934691). No significant changes were seen. One way ANOVA was used for analysis. **B)** Several genes related to hypertrophism were evaluated and showed no change in expression. Expression was measured relative to *Ppia* and graphs are normalized to LFD control expression. One way ANOVA was used for analysis.

## Notes

### Competing Interest Statement

The authors have declared no competing interest.

## References

Aihara, Y., Kurabayashi, M., Saito, Y., Ohyama, Y., Tanaka, T., Takeda, S., Tomaru, K., Sekiguchi, K., Arai, M., Nakamura, T. & Nagai, R. 2000. Cardiac ankyrin repeat protein is a novel marker of cardiac hypertrophy: role of M-CAT element within the promoter. Hypertension, 36, 48–53.

Almon, R. R., Dubois, D. C., Lai, W., Xue, B., Nie, J. & Jusko, W. J. 2009. Gene expression analysis of hepatic roles in cause and development of diabetes in Goto-Kakizaki rats. J Endocrinol, 200, 331–46.

Andersson, U., Filipsson, K., Abbott, C. R., Woods, A., Smith, K., Bloom, S. R., Carling, D. & Small, C. J. 2004. AMP-activated protein kinase plays a role in the control of food intake. J Biol Chem, 279, 12005–8.

Boehringer-Ingelheim. 2021 *AMPK Activator | BI-*9774 [Online]. Available: https://www.opnme.com/molecules/ampk-bi-9774 [Accessed].

Bricogne G. B. E., Brandl M, Flensburg C, Keller P, Paciorek W, et al BUSTER v.2.11.6. ed.

Buque, X., Martinez, M. J., Cano, A., MiquilenA-Colina, M. E., GARCIA-Monzon, C., Aspichueta, P. & Ochoa, B. 2010. A subset of dysregulated metabolic and survival genes is associated with severity of hepatic steatosis in obese Zucker rats. J Lipid Res, 51, 500–13.

Calabrese, M. F., Rajamohan, F., Harris, M. S., Caspers, N. L., Magyar, R., Withka, J. M., Wang, H., Borzilleri, K. A., Sahasrabudhe, P. V., Hoth, L. R., Geoghegan, K. F., Han, S., Brown, J., Subashi, T. A., Reyes, A. R., Frisbie, R. K., Ward, J., Miller, R. A., Landro, J. A., Londregan, A. T., Carpino, P. A., Cabral, S., Smith, A. C., Conn, E. L., Cameron, K. O., Qiu, X. & Kurumbail, R. G. 2014. Structural basis for AMPK activation: natural and synthetic ligands regulate kinase activity from opposite poles by different molecular mechanisms. Structure, 22, 1161–1172.

Carling, D. 2017. AMPK signalling in health and disease. Curr Opin Cell Biol, 45, 31–37.

Chistiakov, D. A., Killingsworth, M. C., Myasoedova, V. A., Orekhov, A. N. & Bobryshev, Y. V. 2017. CD68/macrosialin: not just a histochemical marker. Lab Invest, 97, 4–13.

Cokorinos, E. C., Delmore, J., Reyes, A. R., Albuquerque, B., Kjobsted, R., Jorgensen, N. O., Tran, J. L., Jatkar, A., Cialdea, K., Esquejo, R. M., Meissen, J., Calabrese, M. F., Cordes, J., Moccia, R., Tess, D., Salatto, C. T., Coskran, T. M., Opsahl, A. C., Flynn, D., Blatnik, M., Li, W., Kindt, E., Foretz, M., Viollet, B., Ward, J., Kurumbail, R. G., Kalgutkar, A. S., Wojtaszewski, J. F. P., Cameron, K. O. & Miller, R. A. 2017. Activation of Skeletal Muscle AMPK Promotes Glucose Disposal and Glucose Lowering in Non-human Primates and Mice. Cell Metab, 25, 1147–1159 e10.

Cool, B., Zinker, B., Chiou, W., Kifle, L., Cao, N., Perham, M., Dickinson, R., Adler, A., Gagne, G., Iyengar, R., Zhao, G., Marsh, K., Kym, P., Jung, P., Camp, H. S. & Frevert, E. 2006. Identification and characterization of a small molecule AMPK activator that treats key components of type 2 diabetes and the metabolic syndrome. Cell Metab, 3, 403–16.

Curtis, M. J., Alexander, S., Cirino, G., Docherty, J. R., George, C. H., Giembycz, M. A., Hoyer, D., Insel, P. A., Izzo, A. A., Ji, Y., Macewan, D. J., Sobey, C. G., Stanford, S. C., Teixeira, M. M., Wonnacott, S. & Ahluwalia, A. 2018. Experimental design and analysis and their reporting II: updated and simplified guidance for authors and peer reviewers. Br J Pharmacol, 175, 987–993.

Debose-Boyd, R. A. 2008. Feedback regulation of cholesterol synthesis: sterol-accelerated ubiquitination and degradation of HMG CoA reductase. Cell Res, 18, 609–21.

Dubaisi, S., Barrett, K. G., Fang, H., GUZMAN-Lepe, J., SOTO-Gutierrez, A., Kocarek, T. A. & Runge-Morris, M. 2018. Regulation of Cytosolic Sulfotransferases in Models of Human Hepatocyte Development. Drug Metab Dispos, 46, 1146–1156.

Emsley, P., Lohkamp, B., Scott, W. G. & Cowtan, K. 2010. Features and development of Coot. Acta Crystallogr D Biol Crystallogr, 66, 486–501.

Esquejo, R. M., Salatto, C. T., Delmore, J., Albuquerque, B., Reyes, A., Shi, Y., Moccia, R., Cokorinos, E., Peloquin, M., Monetti, M., Barricklow, J., Bollinger, E., Smith, B. K., Day, E. A., Nguyen, C., Geoghegan, K. F., Kreeger, J. M., Opsahl, A., Ward, J., Kalgutkar A. S., Tess, D., Butler, L., Shirai, N., Osborne, T. F., Steinberg, G. R., Birnbaum, M. J., Cameron, K. O. & Miller, R. A. 2018. Activation of Liver AMPK with PF-06409577 Corrects NAFLD and Lowers Cholesterol in Rodent and Primate Preclinical Models. EBioMedicine, 31, 122–132.

Evans, P. R. & Murshudov, G. N. 2013. How good are my data and what is the resolution? Acta Crystallogr D Biol Crystallogr, 69, 1204–14.

Garcia, D., Hellberg, K., Chaix, A., Wallace, M., Herzig, S., Badur, M. G., Lin, T., Shokhirev, M. N., Pinto, A. F. M., Ross, D. S., Saghatelian, A., Panda, S., Dow, L. E., Metallo, C. M. & Shaw, R. J. 2019. Genetic Liver-Specific AMPK Activation Protects against Diet-Induced Obesity and NAFLD. Cell Rep, 26, 192–208 e6.

Gluais-Dagorn, P., Foretz, M., Steinberg, G. R., Batchuluun, B., Zawistowska-Deniziak, A., Lambooij, J. M., Guigas, B., Carling, D., Monternier, P. A., Moller, D. E., Bolze, S. & Hallakou-Bozec, S. 2022. Direct AMPK Activation Corrects NASH in Rodents Through Metabolic Effects and Direct Action on Inflammation and Fibrogenesis. Hepatol Commun, 6, 101–119.

Goedeke, L., Bates, J., Vatner, D. F., Perry, R. J., Wang, T., Ramirez, R., Li, L., Ellis, M. W., Zhang, D., Wong, K. E., Beysen, C., Cline, G. W., Ray, A. S. & Shulman, G. I. 2018. Acetyl-CoA Carboxylase Inhibition Reverses NAFLD and Hepatic Insulin Resistance but Promotes Hypertriglyceridemia in Rodents. Hepatology, 68, 2197–2211.

Grahame Hardie, D. 2014. AMP-activated protein kinase: a key regulator of energy balance with many roles in human disease. J Intern Med, 276, 543–59.

Hansen, H. H., Feigh, M., Veidal, S. S., Rigbolt, K. T., Vrang, N. & Fosgerau, K. 2017. Mouse models of nonalcoholic steatohepatitis in preclinical drug development. Drug Discov Today, 22, 1707–1718.

Himmelsbach F. L. E., Wagner H. 2015. WIPO Patent WO2015007669.

Hu, L., Su, L., Dong, Z., Wu, Y., Lv, Y., George, J. & Wang, J. 2019. AMPK agonist AICAR ameliorates portal hypertension and liver cirrhosis via NO pathway in the BDL rat model. J Mol Med (Berl), 97, 423–434.

Huang, X., Li, Z., Hu, J., Yang, Z., Liu, Z., Zhang, T., Zhang, C. & Yuan, B. 2019. Knockout of Wdr1 results in cardiac hypertrophy and impaired cardiac function in adult mouse heart. Gene, 697, 40–47.

Jouihan, H., Will, S., Guionaud, S., Boland, M. L., Oldham, S., Ravn, P., Celeste, A. & Trevaskis, J. L. 2017. Superior reductions in hepatic steatosis and fibrosis with co-administration of a glucagon-like peptide-1 receptor agonist and obeticholic acid in mice. Mol Metab, 6, 1360–1370.

Kim, C. W., Addy, C., Kusunoki, J., Anderson, N. N., Deja, S., Fu, X., Burgess, S. C., Li, C., Ruddy, M., Chakravarthy, M., Previs, S., Milstein, S., Fitzgerald, K., Kelley, D. E. & Horton, J. D. 2017. Acetyl CoA Carboxylase Inhibition Reduces Hepatic Steatosis but Elevates Plasma Triglycerides in Mice and Humans: A Bedside to Bench Investigation. Cell Metab, 26, 394–406 e6.

Kim, M., Hunter, R. W., Garcia-Menendez, L., Gong, G., Yang, Y. Y., Kolwicz, S. C.J.R. Xu, J., Sakamoto, K., Wang, W. & Tian, R. 2014. Mutation in the gamma2-subunit of AMP-activated protein kinase stimulates cardiomyocyte proliferation and hypertrophy independent of glycogen storage. Circ Res, 114, 966–75.

Konerman, M. A., Jones, J. C. & Harrison, S. A. 2018. Pharmacotherapy for NASH: Current and emerging. J Hepatol, 68, 362–375.

Liang, H. & Ward, W. F. 2006. PGC-1alpha: a key regulator of energy metabolism. Adv Physiol Educ, 30, 145–51.

Lund, E. G., Guileyardo, J. M. & Russell, D. W. 1999. cDNA cloning of cholesterol 24-hydroxylase, a mediator of cholesterol homeostasis in the brain. Proc Natl Acad Sci U S A, 96, 7238–43.

Mast, N., Norcross, R., Andersson, U., Shou, M., Nakayama, K., Bjorkhem, I. & Pikuleva, I. A. 2003. Broad substrate specificity of human cytochrome P450 46A1 which initiates cholesterol degradation in the brain. Biochemistry, 42, 14284–92.

Mccoy, A. J., Grosse-Kunstleve, R. W., Adams, P. D., Winn, M. D., Storoni, L. C. & Read, R. J. 2007. Phaser crystallographic software. J Appl Crystallogr, 40, 658–674.

Molkentin, J. D. 2004. Calcineurin-NFAT signaling regulates the cardiac hypertrophic response in coordination with the MAPKs. Cardiovasc Res, 63, 467–75.

Myers, R. W., Guan, H. P., Ehrhart, J., Petrov, A., Prahalada, S., Tozzo, E., Yang, X., Kurtz, M. M., Trujillo, M., Gonzalez Trotter, D., Feng, D., Xu, S., Eiermann, G., Holahan, M. A., Rubins, D., Conarello, S., Niu, X., Souza, S. C., Miller, C., Liu, J., Lu, K., Feng, W., Li, Y., Painter, R. E., Milligan, J. A., He, H., Liu, F., Ogawa, A., Wisniewski, D., Rohm, R. J., Wang, L., Bunzel, M., Qian, Y., Zhu, W., Wang, H., Bennet, B., Lafranco Scheuch, L., Fernandez, G. E., Li, C., Klimas, M., Zhou, G., VAN Heek, M., Biftu, T., Weber, A., Kelley, D. E., Thornberry, N., Erion, M. D., Kemp, D. M. & Sebhat, I. K.. 2017. Systemic pan-AMPK activator MK-8722 improves glucose homeostasis but induces cardiac hypertrophy. Science, 357, 507–511.

Oldham, S., Rivera, C., Boland, M. L. & Trevaskis, J. L. 2019. Incorporation of a Survivable Liver Biopsy Procedure in Mice to Assess Non-alcoholic Steatohepatitis (NASH) Resolution. J Vis Exp.

Olivier, S., Foretz, M. & Viollet, B. 2018. Promise and challenges for direct small molecule AMPK activators. Biochem Pharmacol, 153, 147–158.

Percie du Sert, N., Hurst, V., Ahluwalia, A., Alam, S. Avey, M. T., Baker, M., Browne, W. J. Clark, A., Cuthill, I. C., Dirnagl, U. Emerson, M. Garner, P. Holgate, S. T. Howells, D. W., Karp, N. A. Lazic, S. E. Lidster, K. Maccallum, C. J., Macleod, M. Pearl, E. J., Petersen, O. H., Rawle, F. Reynolds, P., Rooney, K. Sena, E. S. Silberberg, S. D. Steckler, T. & Wurbel, H. 2020. The ARRIVE guidelines 2.0: Updated guidelines for reporting animal research. PLoS Biol, 18, e3000410.

Pollard, A. E., Martins, L., Muckett, P. J., Khadayate, S., Bornot, A., Clausen, M., Admyre, T., Bjursell, M., Fiadeiro, R., Wilson, L., Whilding, C., Kotiadis, V. N., Duchen, M. R., Sutton, D., Penfold, L., Sardini, A., Bohlooly, Y. M., Smith, D. M., Read, J. A., Snowden, M. A., Woods, A. & Carling, D. 2019. AMPK activation protects against diet induced obesity through Ucp1-independent thermogenesis in subcutaneous white adipose tissue. Nat Metab, 1, 340–349.

R_CORE_TEAM 2020. R: A language and environment for statistical computing. R Foundation for Statistical Computing. Vienna, Austria.

Santolini, M., Romay, M. C., Yukhtman, C. L., Rau, C. D., Ren, S., Saucerman, J. J., Wang, J. J., Weiss, J. N., Wang, Y., Lusis, A. J. & Karma, A. 2018. A personalized, multiomics approach identifies genes involved in cardiac hypertrophy and heart failure. NPJ Syst Biol Appl, 4, 12.

Smart Os, W. T., Sharff A, Flensburg C, Keller P, Paciorek W, et al. 2011. Grade. 1.2.9 ed.: Global Phasing Ltd.

Song, W., Wang, H. & Wu, Q. 2015. Atrial natriuretic peptide in cardiovascular biology and disease (NPPA). Gene, 569, 1–6.

Soto-Acosta, R. Bautista-Carbajal, P., Cervantes-Salazar, M. Angel-Ambrocio, A. H. & Del Angel, R. M. 2017. DENV up-regulates the HMG-CoA reductase activity through the impairment of AMPK phosphorylation: A potential antiviral target. PLoS Pathog, 13, e1006257.

Stefanovic, L. & Stefanovic, B. 2006. Mechanism of direct hepatotoxic effect of KC chemokine: sequential activation of gene expression and progression from inflammation to necrosis. J Interferon Cytokine Res, 26, 760–70.

Steinberg, G. R., Macaulay, S. L., Febbraio, M. A. & Kemp, B. E. 2006. AMP-activated protein kinase--the fat controller of the energy railroad. Can J Physiol Pharmacol, 84, 655–65.

Sun, C. 2016. Effect of fasting time on measuring mouse blood glucose level. Int J Clin Exp Med, 9, 4186–4189

Trevaskis, J. L., Griffin, P. S., Wittmer, C., Neuschwander-Tetri, B. A., Brunt, E. M. Dolman, C. S. Erickson, M. R., Napora, J. Parkes, D. G. & Roth, J. D. 2012. Glucagon-like peptide-1 receptor agonism improves metabolic, biochemical, and histopathological indices of nonalcoholic steatohepatitis in mice. Am J Physiol Gastrointest Liver Physiol, 302, G762–72.

Williams, C. D., Stengel, J., Asike, M. I., Torres, D. M., Shaw, J., Contreras, M., Landt, C. L. & Harrison, S. A. 2011. Prevalence of nonalcoholic fatty liver disease and nonalcoholic steatohepatitis among a largely middle-aged population utilizing ultrasound and liver biopsy: a prospective study. Gastroenterology, 140, 124–31.

Winter, G., Waterman, D. G., Parkhurst, J. M., Brewster, A. S., Gildea, R. J., Gerstel, M., Fuentes-Montero, L. Vollmar, M. Michels-Clark, T. Young, I. D. Sauter, N. K. & Evans, G. 2018. DIALS: implementation and evaluation of a new integration package. Acta Crystallogr D Struct Biol, 74, 85–97.

Woods, A., Williams, J. R., Muckett, P. J., Mayer, F. V., Liljevald, M., Bohlooly, Y. M. & Carling, D. 2017. Liver-Specific Activation of AMPK Prevents Steatosis on a High-Fructose Diet. Cell Rep, 18, 3043–3051.

Wu, X., Zhang, J., Ge, H., Gupte, J., Baribault, H., Lee, K. J., Lemon, B., Coberly, S., Gong, Y., Pan, Z., Rulifson, I. C., Gardner, J., Richards, W. G. & Li, Y. 2015. Soluble CLEC2 Extracellular Domain Improves Glucose and Lipid Homeostasis by Regulating Liver Kupffer Cell Polarization. EBioMedicine, 2, 214–24.

Xiao, B., Sanders, M. J., Carmena, D., Bright, N. J., Haire, L. F., Underwood, E., Patel, B. R., Heath, R. B., Walker, P. A., Hallen, S., Giordanetto, F., Martin, S. R., Carling, D. & Gamblin, S. J. 2013. Structural basis of AMPK regulation by small molecule activators. Nat Commun, 4, 3017.

Yang, X., Mudgett, J., Bou-About, G. Champy, M. F. Jacobs, H. Monassier, L. Pavlovic, G. Sorg, T. Herault, Y., Petit-Demouliere, B. Lu, K. Feng, W. Wang, H. Ma, L. J. Askew, R. Erion, M. D. Kelley, D. E. Myers, R. W. Li, C. & Guan, H. P. 2016. Physiological Expression of AMPKgamma2RG Mutation Causes Wolff-Parkinson-White Syndrome and Induces Kidney Injury in Mice. J Biol Chem, 291, 23428–23439.

Younossi, Z. M., Koenig, A. B., Abdelatif, D., Fazel, Y., Henry, L. & Wymer, M. 2016. Global epidemiology of nonalcoholic fatty liver disease-Meta-analytic assessment of prevalence, incidence, and outcomes. Hepatology, 64, 73–84.

